# Mapping Lesions that Cause Psychosis to a Human Brain Circuit and Proposed Stimulation Target

**DOI:** 10.1101/2024.04.28.591471

**Authors:** Andrew R. Pines, Summer B. Frandsen, William Drew, Garance M. Meyer, Calvin Howard, Stephan T. Palm, Frederic L.W.V.J. Schaper, Christopher Lin, Konstantin Butenko, Michael A. Ferguson, Maximilian U. Friedrich, Jordan H. Grafman, Ari D. Kappel, Clemens Neudorfer, Natalia S. Rost, Lauren L. Sanderson, Joseph J. Taylor, Ona Wu, Isaiah Kletenik, Jacob W. Vogel, Alexander L. Cohen, Andreas Horn, Michael D. Fox, David Silbersweig, Shan H. Siddiqi

## Abstract

**Importance:** Identifying anatomy causally involved in psychosis could inform therapeutic neuromodulation targets for schizophrenia.

**Objective:** To assess whether lesions that cause secondary psychosis have functional connections to a common brain circuit.

**Design:** This case-control study mapped functional connections of published cases of lesions causing secondary psychosis compared with control lesions unassociated with psychosis.

**Setting:** This study was conducted in a computational laboratory.

**Participants:** Published cases of lesion-induced psychosis were analyzed. Included subjects had documented brain lesions associated with new-onset psychotic symptoms without prior history of psychosis. Control cases included 1156 patients with lesions not associated with psychosis. Generalizability across lesional datasets was assessed using an independent cohort of 181 patients with brain lesions who subsequently underwent neurobehavioral testing.

**Exposures:** Lesions causing secondary psychosis.

**Main Outcomes and Measures:** Psychosis or no psychosis.

**Results:** 153 lesions from published cases were determined to be causal of psychosis (65 [42%] male; mean [SD] age, 50.0 [20.8] years), 42 of which were described as “schizophrenia” or “schizophrenia-like”. Lesions that caused secondary psychosis mapped to a common brain circuit defined by functional connectivity to the posterior subiculum of the hippocampus (84% functional overlap, p_FWE_<5 x 10^-5^). At a lower statistical threshold (75%> overlap, p_FWE_<5 x 10^-4^), this circuit included the ventral tegmental area, retrosplenial cortex, lobule IX and dentate nucleus of the cerebellum, and the mediodorsal and midline nuclei of the thalamus. This circuit was consistent when derived from “schizophrenia-like” cases (spatial r=0.98). We repeated these analyses after excluding lesions intersecting the hippocampus (n=47) and found a consistent functional connectivity profile (spatial r=0.98) with the posterior subiculum remaining the center of connectivity (>75% overlap, p_FWE_<5x10^-5^), demonstrating a circuit-level effect. In an independent observational cohort of patients with penetrating head trauma (n=181), lesions associated with symptoms of psychosis exhibited significantly similar connectivity profiles to the lesion-derived psychosis circuit (suspiciousness, p=0.025; unusual thought content, p=0.046). Voxels in the rostromedial prefrontal cortex (rmPFC) are highly correlated with this psychosis circuit (spatial r=0.82), suggesting the rmPFC as a promising TMS target for psychosis.

**Conclusions and Relevance:** Lesions that cause secondary psychosis affect a common brain circuit in the hippocampus. These results can help inform therapeutic neuromodulation targeting.

**Key Points:** *Question:* Do lesions that cause psychosis affect a common brain circuit?

*Findings:* This case-control study found that lesions causing psychosis specifically affected a common functional circuit aligning with the posterior subiculum of the hippocampus. This functional circuit was consistent across different psychotic symptoms, suggesting a shared neural substrate for psychosis. A similar circuit was derived when excluding lesions directly intersecting the hippocampus, indicating a circuit-level effect.

*Meaning:* Identifying a common brain circuit causally involved in psychotic symptoms suggests that the hippocampus may be pivotal in the pathophysiology and treatment targeting of psychotic disorders.

## Introduction

Symptoms of psychosis — delusions, hallucinations, and disorganized thought — are a significant contributor to global disease burden^1^. Current pharmacological treatments can be incompletely effective and often have intolerable side effects^2,3^. Identifying neuroanatomy causally involved in psychosis and determining its functional connections could guide targeting of therapeutic brain stimulation^4^.

Neuroanatomy of psychosis can be investigated with neuroimaging correlates in patients with schizophrenia, a disease defined by chronic primary psychosis. Functional neuroimaging studies have reported global dysconnectivity as well as aberrancy in default mode and frontoparietal networks^5–8^, while structural neuroimaging studies have shown abnormalities in the corpus callosum and limbic network^9^ along with grey-matter reductions throughout the cortex, most consistently in the medial temporal lobes^10,11^. While these correlates provide valuable insights into the disease process of schizophrenia, observed abnormalities could be spurious, epiphenomenal, or compensatory processes^10,12^.

Lesion studies can complement these correlates by informing the direction of causality^13^. This is particularly important for therapeutic targeting, as modulating a compensatory process could exacerbate a symptom rather than improve it. Lesions that cause psychosis, however, can occur in disparate brain regions^14^. A possible explanation is that the anatomical unit responsible for psychosis is not a single brain region, but connections between multiple brain regions. This hypothesis can be investigated using a normative brain functional connectome to determine if lesions in disparate locations across the brain affect a common functional circuit^15^.

Causal inferences from lesion studies have guided therapeutic targeting for multiple neuropsychiatric disorders. For instance, the most effective TMS targets for depression are functionally connected to lesions that cause depression^16–19^. Similarly, lesion-derived circuits have revealed TMS targets for addiction, pain, and other syndromes^16,20,21^, suggesting that a lesion-derived psychosis circuit may be a reasonable TMS target.

Here, we investigated lesions causing symptoms that mimic schizophrenia. We derived and cross-validated a circuit based on the functional connectivity of published cases of lesion-induced psychosis. Finally, we performed an exploratory analysis using this lesion-derived circuit to identify a novel target for future Transcranial Magnetic Stimulation (TMS) studies in psychosis.

## Methods

This study used publicly available data and did not involve direct human subjects research. Consequently, it was determined exempt from institutional review board oversight by Brigham and Women’s Hospital, Boston, Massachusetts and from obtaining informed consent based on the secondary use of public data. . For full details on the dataset and each analysis, see eMethods in Supplement 1.

### Case Selection

We systematically searched PubMed according to PRISMA Guidelines^22^ to identify published cases of brain lesions associated with psychosis (eFigure 1 in Supplement 1). Cases were included if the lesion was radiologically determined to be acute and symptoms developed within 1 year, the patient had negative imaging within a year before psychosis began, or resolution of the lesion coincided with symptom resolution. Cases with a prior history of psychosis were excluded.

**Figure 1.**
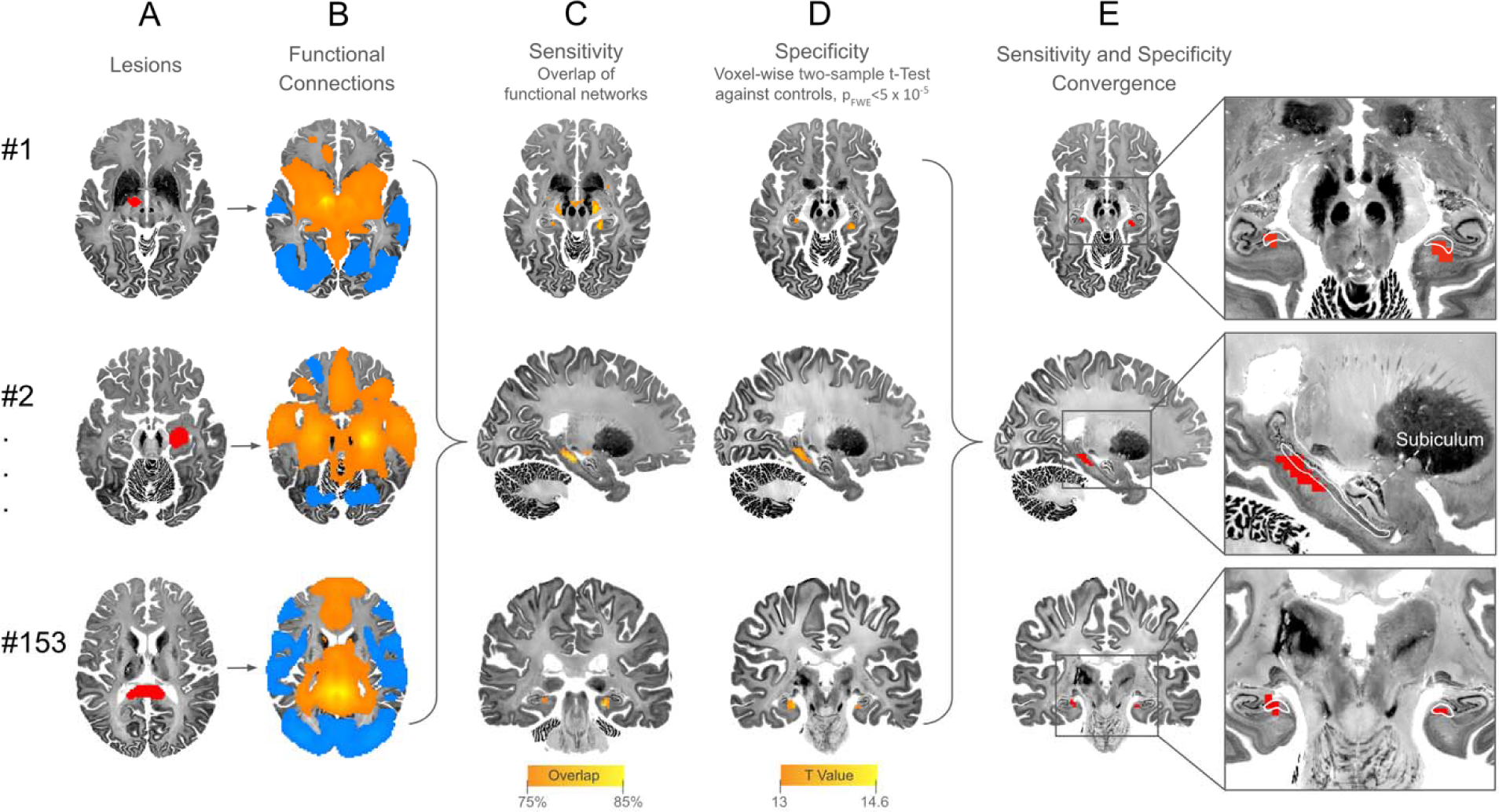
Lesions that Cause Psychosis Map to a Common Circuit of Functional Connectivity **(a)** Each lesion traced manually onto a standard brain atlas^73^. **(b)** Normative connectivity of each lesion was determined using a normative connectome database (n=1000)^24^. **(c)** Sensitivity analysis showing the overlap of functional connectivity of >75% of lesions, thresholded at |T|>7. **(d)** Specificity analysis showing the results of a two-sample t-test (p_FWE_<5 x 10^-5^) comparing functional connectivity of lesions that cause psychosis and control lesions. **(e)** Convergence map depicting the posterior subiculum as the most sensitive and specific region affected by lesions that cause psychosis. Subiculum is defined by the CoBrALab Merged Atlas^74^.

### Lesion Network Mapping

Lesions were reconstructed by manually tracing published images of lesions onto an MNI template using 3D Slicer (Boston, MA; https://www.slicer.org/)23 (Figure1a; eFigure 2 in Supplement 1). The functional connectivity of each lesion was estimated by correlating the lesion location with a normative functional connectome of healthy adult participants (n=1000)^24^, generating a brain-wide functional connectivity map for each lesion (Figure 1B).

**Figure 2.**
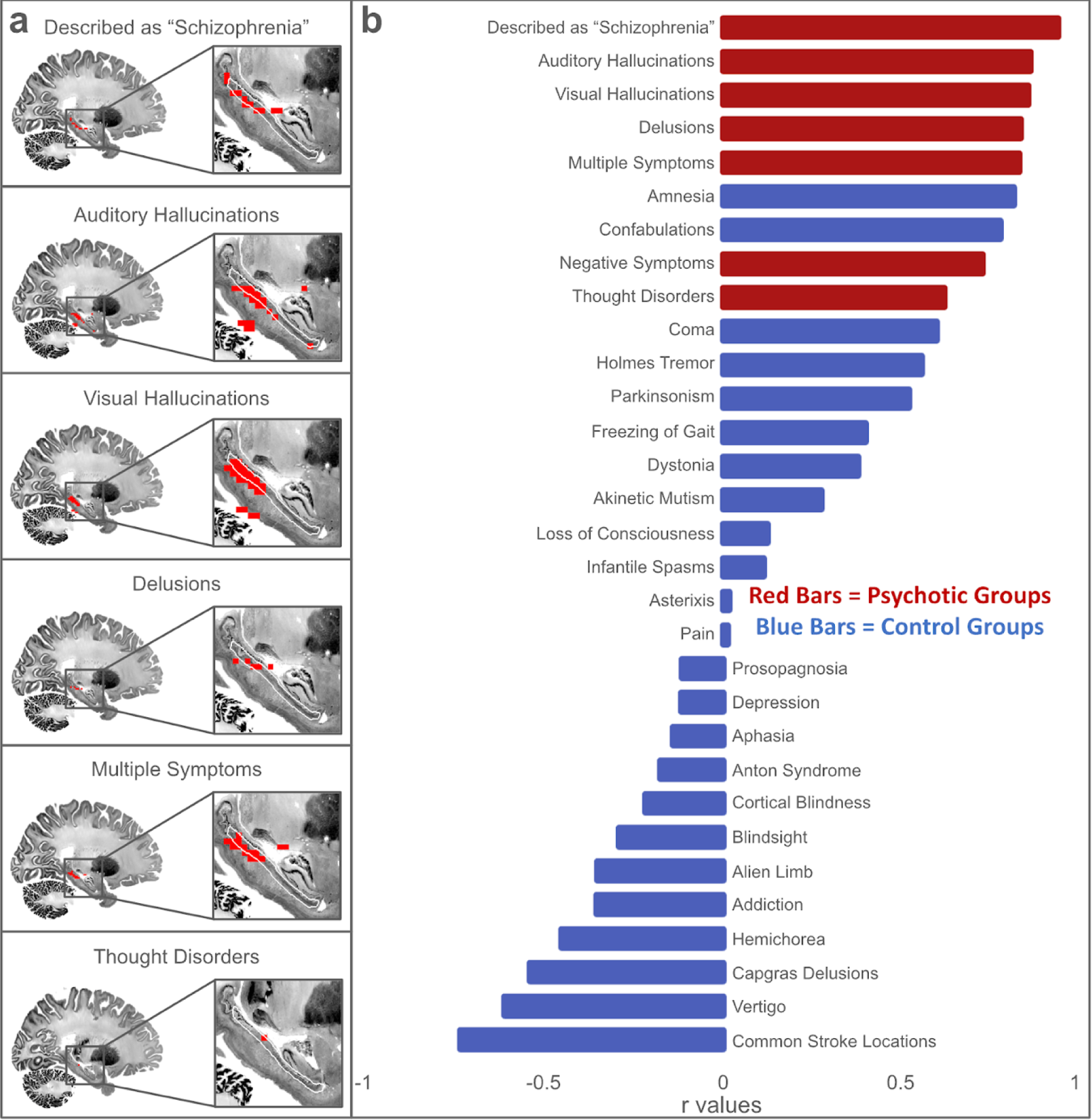
Functional connections of lesions that cause different symptoms of psychosis share a common functional connection to the posterior subiculum of the hippocampus, and are more correlated with each other than with lesions that cause other symptoms **a)** Each grouping of psychotic symptoms had peak sensitivity (overlap >75%) and specificity (p_FWE_<5 x 10^-4^) in the posterior subiculum of the hippocampus (outlined in white). No individual voxels from the Negative Symptoms subgroup remained significant after correcting for multiple comparisons. **b)** Spatial correlation values between the psychosis circuit and each symptom group. When correlating between the psychosis circuit and each psychotic symptom group, lesions from the psychotic group being examined were excluded from the psychosis circuit they were being correlated to.

First, in a sensitivity analysis, we identified common voxels functionally connected to lesions causing psychosis. Each map was thresholded |t| ≥ 7 (Figure 1C) as in prior work^25^. To ensure results were not threshold-dependent, analyses were repeated with alternative thresholds 5 and 10, and further thresholded with a one-sample t-test.

Next, in a specificity analysis, we compared functional connectivity maps of lesions causing psychosis with control lesions not associated with psychosis (n=1156) (Figure 1D). We used Permutation Analysis of Linear Models (PALM) (https://fsl.fmrib.ox.ac.uk/fsl/fslwiki/PALM)^26^ to correct for multiple comparisons across voxels.

Voxels that passed both tests were considered sensitive and specific to functional connections of lesions that cause psychosis (Figure 1E).

### Analysis of different symptoms of psychosis

To determine if different psychotic symptoms mapped to similar anatomy, we categorized lesions by their resulting symptom(s): delusions, thought disorders, auditory hallucinations, visual hallucinations, and negative symptoms. We derived a sensitivity and specificity map for each symptom group by comparing the functional connectivity of that symptom group with the control group in a voxel-wise two-sample t-test, as above. We then performed a leave-one-symptom-out analysis using spatial correlations to compare the specificity map of each symptom group with a specificity map generated from the remaining psychotic symptom groups. For comparators, we repeated this analysis for each symptom caused by control lesions (Figure 2A).

### Analysis of different symptoms of psychosis presenting in isolation

The above analyses used cases in which multiple psychotic symptoms co-occur to model schizophrenia. To determine if lesions causing a single psychotic symptom mapped to distinct anatomy, we repeated the lesion network mapping analysis including only lesions that caused each psychotic symptom in isolation. In addition to FWE correction across voxels, we used False Discovery Rate correction to account for analysis across multiple symptom groupings.

### Independent validation

We used the Vietnam Head Injury Study (VHIS)^27^ for external validation. This dataset has lesions from 181 male American military Veterans (mean age 58.4 ± 3.1) who completed the neurobehavioral rating scale (NBRS) after penetrating brain trauma. We estimated functional connectivity of each lesion using the same normative functional connectivity database as the previous analyses. We correlated the functional maps of the VHIS lesions with their associated behavioral ratings using a voxel-wise Pearson correlation. The resulting symptom-level maps were compared to the psychosis circuit produced from the specificity analysis above, yielding a correlation between symptom-level maps in an independent dataset and the psychosis specificity map. We hypothesized that the psychosis circuit would be preferentially connected to lesions that induce psychosis-like symptoms in this dataset - i.e., “suspiciousness” and “unusual thought content” - relative to 24 control symptoms unrelated to psychosis.

### Identification of therapeutic TMS target

To identify a therapeutic target whose intrinsic functional connectivity is most similar to the psychosis circuit, we estimated the whole-brain connectivity of each voxel using the same normative functional connectome employed in earlier steps (Figure 5a). We then used spatial correlations to compare the connectivity map of each voxel with the psychosis specificity map (Figure 5b). This produced a map denoting voxels whose connectivity best matches the psychosis circuit. To determine the extent to which this TMS target would overlap with the electrical field generated by a magnetic coil, we modeled the electrical field (E-field) of a MagVenture B65 (MagVenture, Farum, Denmark) coil in MNI space using SimNIBS^28^.

### Statistical Analysis

Statistical analyses were performed using MATLAB R2022b (Mathworks, Natick MA). Data were analyzed from June 2022 - April 2024. Methods of type-I error control for each test are listed in eTable 2 in Supplement 1. Non-parametric testing was performed for each instance of a parametric test (eMethods in Supplement 1). More stringent statistical thresholds were used to highlight the most robust findings (Figure Legends).

## Results

### Case Selection

We identified 153 cases of lesions that caused secondary psychosis, including 83 with visual hallucinations, 81 auditory hallucinations, 67 delusions, 25 thought disorder, and 13 with negative symptoms. 87 cases presented with more than one symptom. 42 reports described symptoms as “schizophrenia” or “schizophrenia-like” (eTable 1 in Supplement 1).

### Lesion Network Mapping

Lesions that caused secondary psychosis were not overrepresented in any specific lobe or vascular territory (eFigure 3 in Supplement 1).

**Figure 3.**
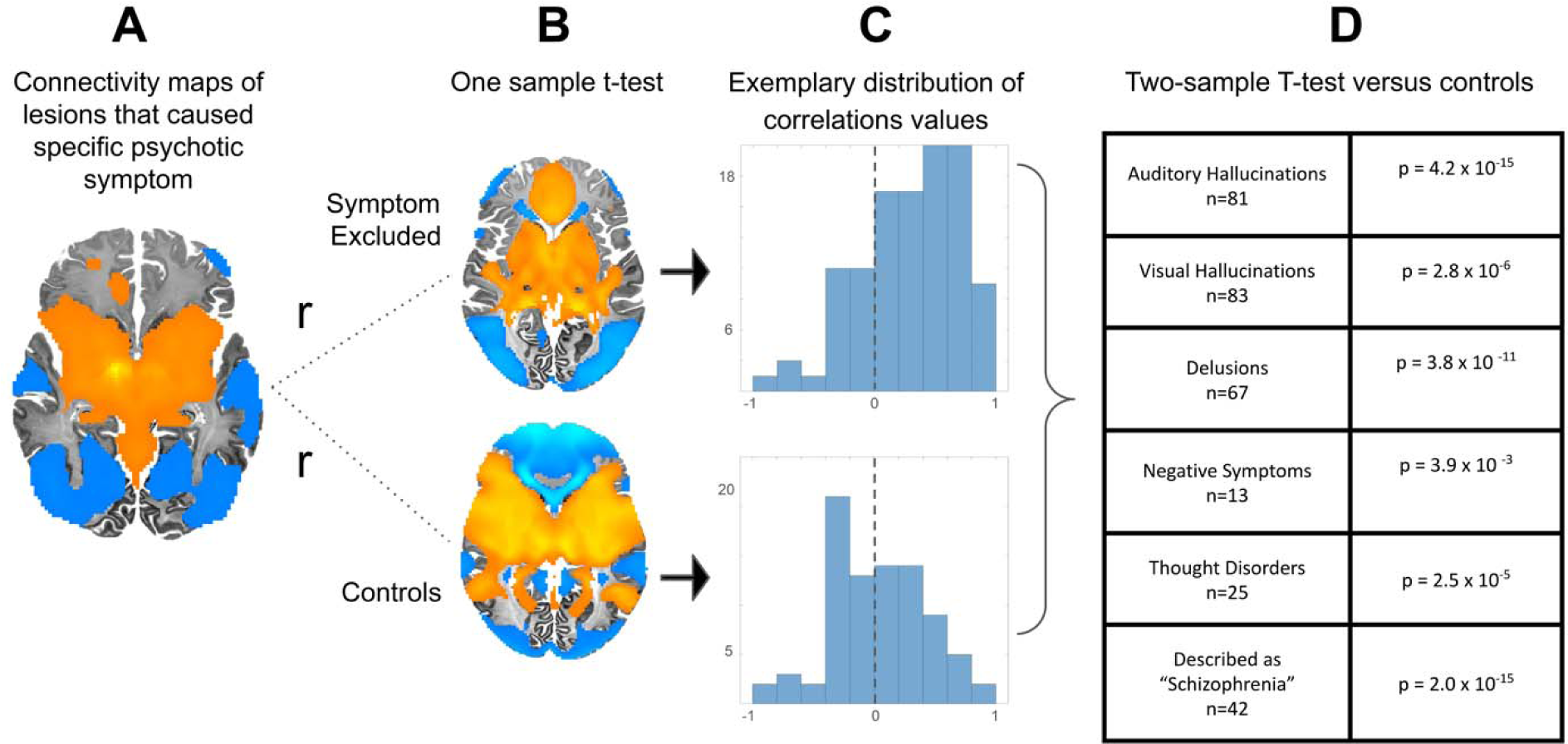
Different symptoms of psychosis map to common circuitry **(a)** Representative example of functional connectivity of a lesion causing a psychotic symptom. **(b)** Connectivity maps of each symptom group were correlated with a map of mean connectivity of lesions that cause psychosis without the symptom being tested and mean connectivity of control lesions. **(c)** Distribution of correlations of each lesion to each of the two comparator groups. Y axis=number of cases; X axis=r value (Range: -1 to 1. The zero value is marked with a dotted line) **(d)** For each group of psychotic symptoms, lesion connectivity was more similar to other psychotic symptoms than to control symptoms.

A sensitivity analysis found that 84% (129/153) of lesions were connected to the posterior subiculum of the hippocampus (Figure 1C). Results were unchanged when using alternative T-value thresholds (spatial r>0.99, peak remained in the subiculum). A one-sample t-test of all 153 maps at each voxel identified the posterior subiculum (p_FWE_<5x10^-4^) as consistently present in functional connectivity maps of psychosis-inducing lesions.

A specificity analysis found the functional connections most specific to lesions that cause secondary psychosis were also located in the posterior subiculum (p_FWE_<5 x 10^-5^) (Figure 1D).

The most sensitive and specific functional connections were in the posterior subiculum (Figure 1E). Secondary peak regions (overlap >75% on sensitivity analysis, p_FWE_<5x10^-4^ on specificity analysis) included the ventral tegmental area, mediodorsal and midline thalamic nuclei, retrosplenial cortex, and lobule IX and dentate nucleus of the cerebellum (eFigure 5 in Supplement 1).

### Lesion Network Mapping without hippocampal lesions

Next, to test whether these results were driven by a circuit-effect or by hippocampal lesions alone, we repeated the analysis after excluding 36 lesions with at least one voxel in the hippocampus. This analysis yielded equal peaks (>75% overlap, p_FWE_<5x10^-5^) in the posterior subiculum of the hippocampus, the ventral tegmental area, mediodorsal and midline nuclei of the thalamus, and lobule IX and dentate gyrus of the cerebellum (eFigure 7 in Supplement 1). The specificity map from this analysis was nearly identical to its counterpart that included hippocampal lesions (spatial r=0.98). Thus, symptoms of psychosis were caused by lesions with functional connectivity to the hippocampus, not just by lesions that intersect the hippocampus.

### Analysis of different symptoms of psychosis

The functional connectivity profiles of most psychotic symptom groups displayed a similar peak in the posterior subiculum (Figure 2a) and were highly similar to each other (median spatial r=0.84, range 0.62-0.94) (Figure 2b). The least similar symptom was thought disorder (r=0.62), which had a primary peak in the corpus callosum and a secondary peak in the subiculum (eFigure 9 in Supplement 1). Of note, this group had a relatively small sample size (n=25).

These results suggest that lesions that cause different symptoms of psychosis have functional connectivity profiles more similar to each other than controls. To further test this hypothesis, we used spatial correlations to compare the functional maps of each symptom group (Figure 3a) to a one-sample t-test of control lesions and a one-sample t-test of psychosis lesions, excluding the symptom group under examination (Figure 3b). We then performed a two-sample t-test between the distributions of correlations with controls and the distributions of correlations with other psychosis-inducing lesions. The functional connectivity of lesions that caused each psychotic symptom was more similar to each other than to control lesions (Figure 3c).

### Analysis of different symptoms of psychosis presenting in isolation

While most lesions caused multiple symptoms, 32 caused isolated visual hallucinations, 24 caused isolated auditory hallucinations, and 8 caused isolated paranoid delusions.

Lesions that cause isolated visual hallucinations localized to the subiculum (eFigure 10 in Supplement 1); however, lesions causing Charles-Bonnet Syndrome by disrupting primary visual input, which would be expected to map to primary visual regions, were excluded from this study because this disease is clearly distinguished from schizophrenia (eMethods in Supplement 1). Lesions that caused isolated auditory hallucinations (see eMethods in Supplement 1 for phenomenological description) were exclusively located in either the brainstem (n=11) or temporal lobe (n=13, seven of which intersected Heschl’s Gyrus). Temporal lobe lesions causing isolated auditory hallucinations had peak connectivity to the superior temporal gyrus (85% overlap, p_FWE_<5x10^-5^) (eFigure 10 in Supplement 1). Brainstem lesions causing isolated auditory hallucinations had peak connectivity to the subcortical auditory system (100% overlap, p_FWE_<5x10^-5^) (eFigure 10 in Supplement 1). Of note, no patients with isolated auditory hallucinations were described as “schizophrenia-like”: hallucinations were simple sounds instead of distinguishable language and patients retained insight that hallucinations were not real (eMethods in Supplement 1). Lesions that caused isolated paranoid delusions localized to the right mediodorsal thalamus (100% overlap, p_FWE_<0.05) (eFigure 10 in Supplement 1).

### Distinguishing from lesions that cause amnesia

Functional connections of lesions that cause secondary psychosis were more similar to functional connections of lesions that cause amnesia than any other control group (spatial r=0.82) (Figure 3A). In a post-hoc analysis, we explored features that distinguish these lesions.

Among the 53 published amnesia cases used to define a memory circuit^29^, six (11%) also had symptoms of psychosis (eMethods in Supplement 1). Among the 153 psychosis cases in this study, 48 reported an acute memory deficit, 7 reported no memory deficit, and 98 did not report any memory testing (eMethods in Supplement 1). To directly contrast psychosis and amnesia, we compared the functional connections of lesions that caused both psychosis and amnesia (n=48) to lesions causing amnesia without psychosis (n=47). Lesions causing psychosis were more functionally connected to aspects of the superior temporal gyrus, the ventral claustrum, and part of the uncinate fasciculus (p_FWE_ < 0.05) (eFigure 11 in Supplement 1). Lesions causing amnesia without psychosis were not associated with any significant voxels.

In an exploratory analysis of laterality, lesions causing psychosis were more likely to involve the right hemisphere (n=9:22:17 Left:Bilateral:Right), while lesions causing amnesia were more likely to involve the left hemisphere (19:20:7) (?^2^ p=0.03).

### Validation in an independent cohort

Lesions from the VHIS with greater connectivity to the psychosis circuit were more likely to be associated with suspiciousness and unusual thought content (p=0.025 and 0.046, respectively) (Figure 4) than the other 24 control symptoms.

**Figure 4.**
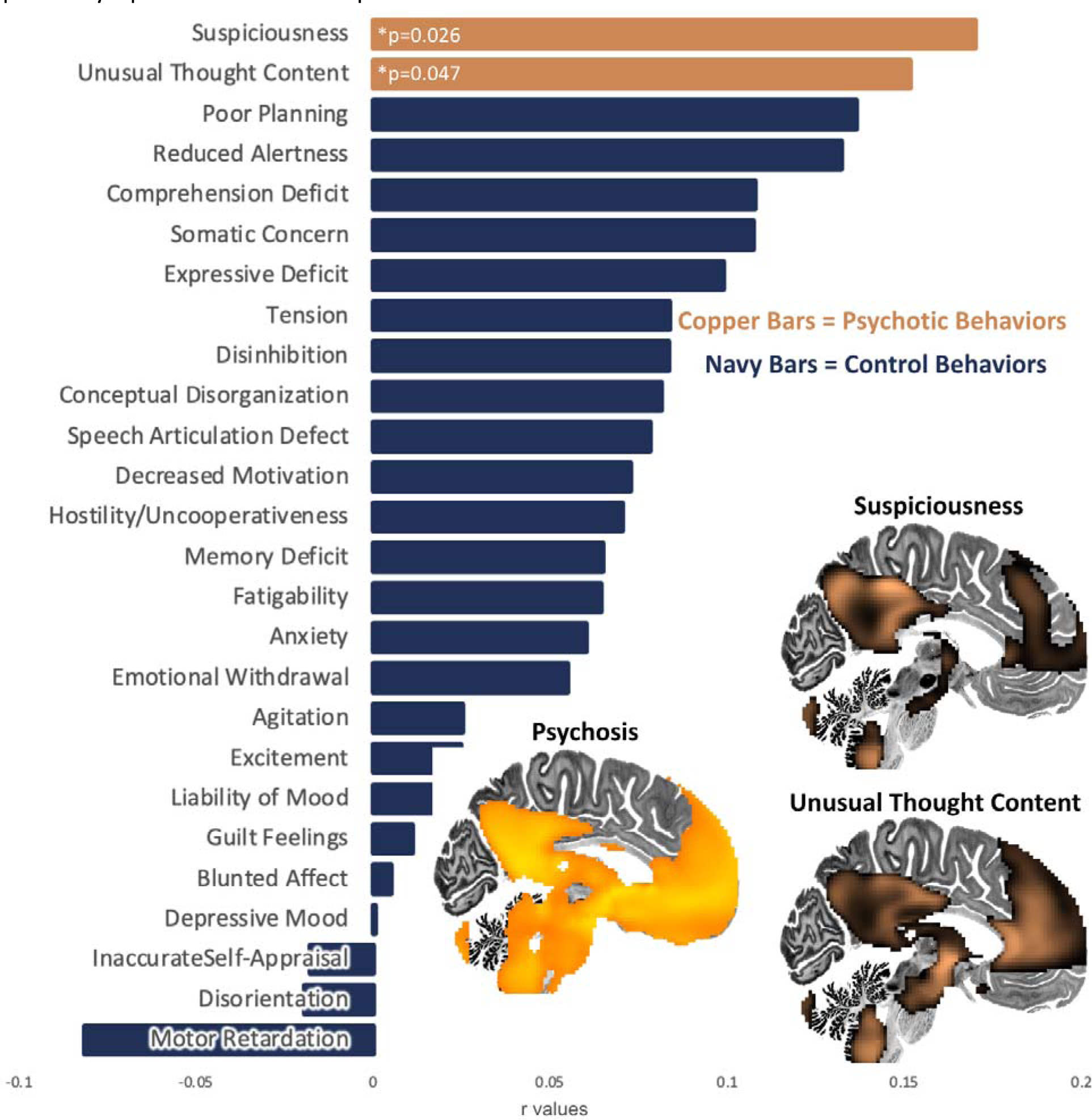
Correlation values between functional maps of lesions causing psychosis and lesions causing specific symptoms from an independent cohort. Functional connectivity maps specific to ‘Suspiciousness’ and ‘Unusual Thought Content’ are shown to illustrate similarity to the psychosis circuit.

**Figure 5.**
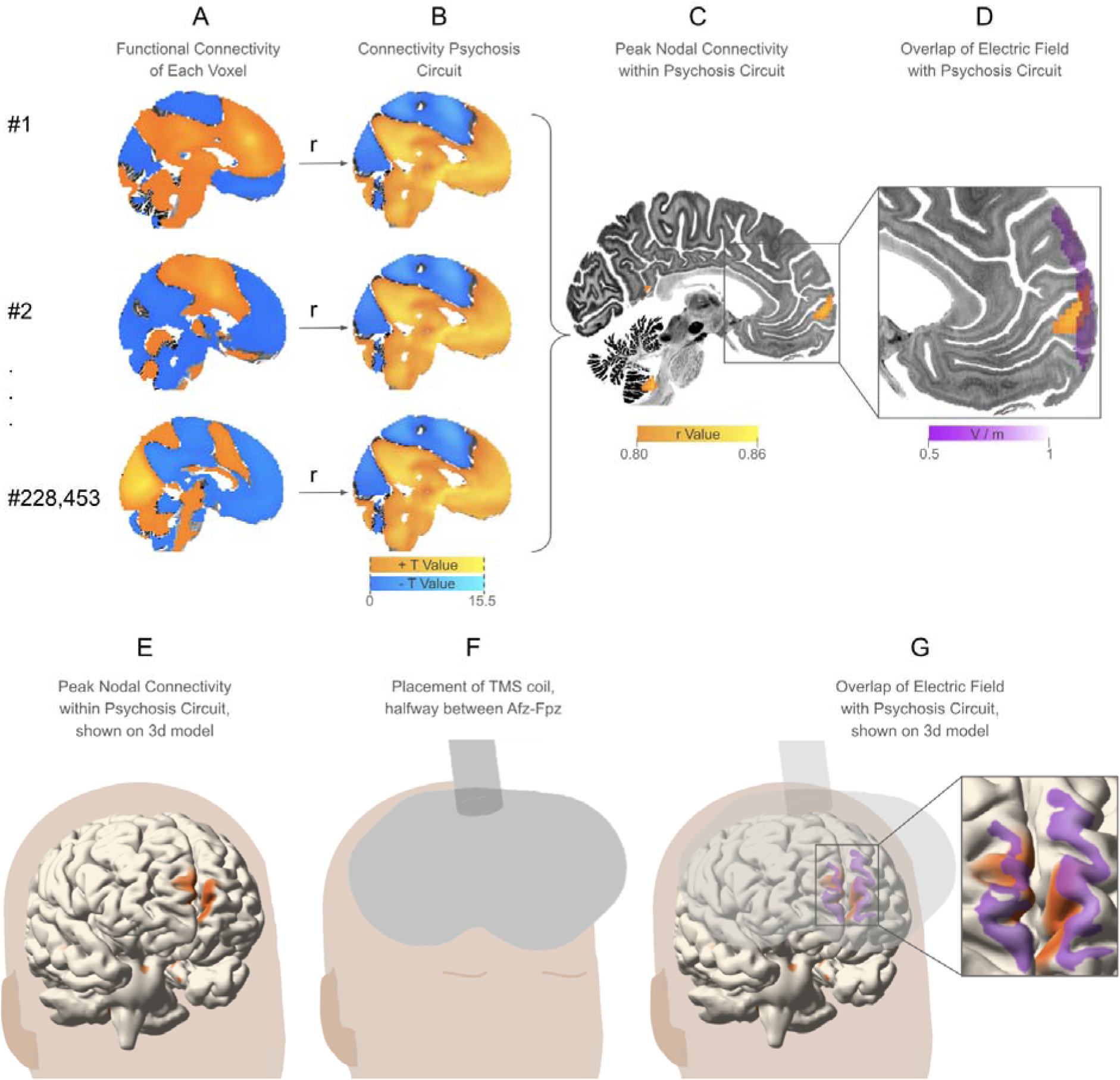
Therapeutic TMS target identified by correlation with each voxel’s intrinsic functional connectivity. **a)** BOLD correlations for each voxel in a 2mm brain template **b)** correlated with unthresholded BOLD correlation for the two-sample t-test between lesions causing psychosis and controls. **c)** identification of voxels with the strongest correlation between intrinsic resting-state BOLD signal and BOLD signal specific to lesions that cause psychosis. **d)** A simulated electric field measured in volts per meter (V/m) overlaps peak nodal connectivity within the psychosis circuit, indicating that the psychosis circuit is sufficiently superficial to be targeted with TMS. **e)** Peak nodal connectivity of the psychosis circuit, identical to panel C, shown on a 3d model^21,76^. **f)** Placement of the TMS coil, halfway between Afz-Fpz. **g)** Magnetic field generated by the TMS coil in the position overlapping peak nodal connectivity within the psychosis circuit, identical to panel D, shown on a 3d model.

### Identification of therapeutic TMS target

The voxel with whole-brain connectivity best matching the psychosis circuit was in the subiculum (spatial r=0.83), but this location is too deep to access with TMS. A site within the right rostromedial prefrontal cortex (rmPFC) (MNI [2,66,2]) was also highly correlated to the psychosis circuit (r=0.82)(Figure 5c,e). This site overlaps with an E-field simulated from a position equidistant between PFz and AFz electrode placements on the 10-20 EEG system, so is sufficiently superficial to serve as a TMS target (Figure 5d,g).

## Discussion

Lesions causing secondary psychosis mapped to a common functional circuit with a peak in the posterior subiculum of the hippocampus. Results were consistent across psychotic symptoms and when excluding lesions that directly touch the hippocampus. This circuit was associated with psychosis-like symptoms in an independent cohort. A region in the rmPFC was highly connected to the circuit and is accessible with TMS. These findings have several important implications.

First, we provide evidence that causal disruption of a hippocampal circuit can produce psychosis. Hippocampal involvement in psychosis is consistent with decades of neuroimaging and lesion studies. To our knowledge, this is the first study to systematically unite these lesion studies into a common circuit-based causal model, as such models have been shown to improve brain stimulation targeting^4,13^. Causal studies have implicated other systems in psychosis, such as N-methyl-D-aspartate receptors (NMDAr), as some NMDAr antagonist drugs and anti-NMDAr encephalitis can produce symptoms closely resembling schizophrenia^30,31^. These symptoms have been hypothesized to result from hippocampal NMDAr modulation^32–34^, but the effect of NMDAr on psychosis has not been definitively localized to a specific brain region. The present study employed spatially-confined lesions that cause secondary psychosis and can therefore be used to infer both the direction of causality and neuroanatomical specificity^13^. This anatomical localization was driven by a circuit-level effect, as a similar circuit was derived even when excluding lesions that touch the hippocampus.

The relationship between the hippocampus and psychosis has been studied extensively (further detailed in Hippocampus and Psychosis Literature in Supplement 1). Atrophy of the hippocampus is one of the most consistent findings on neuroimaging^10,35^. Though patients with schizophrenia do not have a decrease in overall neurons^36,37^, they appear to have fewer interneurons across subfields of the hippocampus^38,39^ and parahippocampal gyrus^40^. Psychotic disorders have been associated with abnormal hippocampal activity^41^, particularly in the anterior CA1 subfield^42–45^, impairing hippocampal recruitment and function^46,47^. This is consistent with our localization to the subiculum, whose primary input is from CA1^48,49^. Our findings may further build on this literature by informing the direction of causality, but our results do not elucidate the mechanism of hippocampal dysfunction, as BOLD signal likely does not resolve cell types, subfields, or structural connections.

The peak region of this circuit is in the posterior subiculum of the hippocampus. Notably, the major afferent and efferent connections of the subiculum include the medial prefrontal cortex^50,51^, along with CA1^48,52^and the retrosplenial^53–55^ and rhinal cortices^56,57^. The subiculum is the primary outflow tract of the hippocampus to the fornix^58,59^, from which the hippocampus communicates with the mamillary bodies, septal nuclei, and anterior nuclei of the thalamus^59–61^. Results from this study are consistent with recent work demonstrating that grey matter atrophy in schizophrenia progresses along structural and functional connections originating from the hippocampus^62,63^.

This hippocampal region was identified across different individual psychotic symptoms, supporting conceptualization of psychosis as a discrete syndrome rather than a constellation of different symptoms. By contrast, lesions causing isolated auditory hallucinations of simple sounds which do not resemble schizophrenia-like symptomatology appear to affect primary auditory regions. The syndrome of schizophrenia affects multiple sensory and cognitive modalities, suggesting that symptoms co-occur as a downstream effect of higher-order dysfunction^64^. This model suggests that the different symptoms of schizophrenia can result from dysfunctional communication between the hippocampus and higher-order processing regions.

Prior work has shown that lesion-derived circuits can reveal TMS targets for addiction, depression, pain, and other syndromes^16,20,21^, suggesting that this lesion-derived psychosis circuit may be a reasonable TMS target. As lesions presumably disrupt communication between regions, we hypothesize that TMS targeted to this circuit should be performed with parameters to increase activity^65^. We identified a target in the rmPFC that is accessible with TMS and is highly connected to our circuit. A similar target has previously been proposed based on task-fMRI studies of reality monitoring in schizophrenia^66^, and TMS to this target improves reality monitoring in healthy controls^67^. A nearby region has also been targeted for addiction and obsessive-compulsive disorders and has been well-tolerated^68^. Sham-controlled TMS trials for schizophrenia have demonstrated reductions in negative symptoms when targeting the cerebellum or dorsolateral prefrontal cortex^69^. Still, no target has consistently improved positive symptoms^70^ (further detailed in Sham-Controlled TMS Trials for Schizophrenia in Supplement 1). Across targets, TMS trials have demonstrated safety in patients with schizophrenia^71^.

## Limitations

There are several limitations to this study. First, lesion-induced psychosis is not schizophrenia, although a similar circuit was derived from lesions causing symptoms described as “schizophrenia-like”. Studying causality in schizophrenia can be challenging due to the difficulty of comprehensively measuring genetic and environmental factors. Future studies may generalize causality of this lesion-derived circuit to primary schizophrenia if modulation of this circuit correlates with therapeutic response.

Next, this study employs the average of a large functional connectome to approximate the connectivity of each lesion. Due to variation in functional connectivity between individuals^72^, it remains unclear if connection impairment occurred in any individual patient. While this may limit inference at the individual patient level, these estimates have previously been shown to be sufficient for brain stimulation target identification^16,20,21^.

While lesion case reports are valuable because they report a stark change that temporally coincides with a focal perturbation, there are also limitations to this approach, such as selection bias and variations in imaging parameters and institutional practices. Case reports are also limited in detail and the standardization of diagnostic assessments. This may differentially impact different symptoms - for instance, thought disorder was often described superficially, thus limiting its characterization. This may explain why the anatomical peaks were slightly different for thought disorder relative to other symptoms, although lesions causing thought disorder still significantly mapped to the subiculum and were still connected to a circuit that is significantly similar to the circuits for other psychotic symptoms.

Lesion studies also carry some inherent limitations: they are observational, rely on imprecise anatomical boundaries, can be confounded by undetected pathological processes, and may oversimplify neurocircuitry into a binary model of active/inactive functioning. Our results do not support this binary model, as lesions functionally connected to the posterior subiculum do not always cause psychosis. Mechanisms of psychosis are undoubtedly influenced by genetics, developmental factors, neurotransmitters, and microcircuitry. While lesions may inform the direction of causality, they do not define a causal mechanism. Mechanistic insights may be gleaned from pharmacological manipulation of relevant receptors, post-mortem gene expression studies, interrogation of neuronal subpopulations, and animal models of neurodevelopment. Our study was not intended to delineate mechanisms of schizophrenia but rather to use a phenocopy of the disease as a model for identifying therapeutic targets.

## Conclusions

This study identified a human brain circuit aligning with the posterior subiculum subfield of the hippocampus that is causally implicated in symptoms of psychosis, particularly in cases that resemble schizophrenia. These findings could guide therapeutic brain stimulation.

## Acknowledgments

There was no funding for this study, and no funding entity or sponsor had a role in the no role in the design and conduct of the study; collection, management, analysis, and interpretation of the data; preparation, review, or approval of the manuscript; and decision to submit the manuscript for publication. MDF has served as a scientific consultant for Magnus Medical and is owner of intellectual property involving the use of functional connectivity to target TMS, which was not used in the present study. SS is an owner of intellectual property involving the use of brain connectivity to target TMS; has served as a scientific consultant for Magnus Medical; has received investigator-initiated research funding from Neuronetics and Brainsway; has received speaking fees from Brainsway and Otsuka (for PsychU.org); and is a shareholder in Brainsway (publicly traded) and Magnus Medical (not publicly traded). All other authors declare they have no competing interests. ARP had full access to all the data in the study and takes responsibility for the integrity of the data and the accuracy of the data analysis.

