## Supplement 1 for "Mapping Lesions that Cause Psychosis to a Human Brain Circuit and Proposed Stimulation Target"

**Supplemental Online Content**

**eMethods**

**eTable 1.** Case references, demographics, and characteristics

**eFigure 1.** Results of PRISMA-formatted literature review

**eFigure 2.** Example Lesions Associated with Psychosis (6 of 153)

**eFigure 3.** Lesions associated with psychosis and control lesions were distributed throughout vascular territories and brain regions.

**eFigure 4.** Voxel-wise two-sample t-test between lesions that cause psychosis and control lesions not associated with psychosis.

**eFigure 5.** Secondary Peak Regions of Psychosis Circuit

**eFigure 6.** Convergence of Sensitivity and Specificity Tests, Accounting for Age and Sex Covariates

**eFigure 7.** Functional Connectivity of Lesions that Cause Psychosis is Consistent When Excluding Hippocampal Lesions

**eFigure 8.** Leave-one-out analysis, leaving out the functional map of each lesion

**eFigure 9.** Lesions that cause thought disorder map to different regions of a similar circuit.

**eFigure 10.** Lesions that cause isolated symptoms of psychosis map to distinct regions.

**eFigure 11.** Voxel-wise two-sample t-tests comparing lesions that cause psychosis with amnesia and lesions that cause only amnesia.

### eMethods

All statistical tests were performed using MATLAB R2022b except as otherwise specified.

**Case Selection**

We identified published cases of brain lesions associated with psychosis with a systematic search on PubMed according to PRISMA Guidelines^1^ (eFigure 1). We excluded cases with a prior history of psychosis. Cases were included only if there was evidence of a clear temporal relationship between the lesion and the psychotic symptoms (the lesion was radiologically determined to be acute on imaging, the patient had negative imaging within the past year before the psychotic symptoms started, or resolution of the lesion coincided with symptom resolution). Symptoms of schizophrenia are commonly split into ‘positive’ symptoms (delusions, thought disorders, hallucinations) and ‘negative’ symptoms (apathy, avolition, cognitive deficits). Cases reporting symptoms of apathy, avolition, and cognitive deficits that presented in isolation rather than as part of a psychotic syndrome were not included in the literature review. Although these can be classified as negative symptoms of schizophrenia, stroke-induced apathy or avolition are well-described clinical phenomena that are usually distinct from psychosis^2^. Cases were not included if psychotic symptoms were accompanied by confusion, disorientation, or inattention, with all symptoms resolving days to weeks after the lesion. These symptoms are better explained by delirium or post-traumatic confusion rather than the lesion location. Cases were included in the category of ‘thought disorder’ if the subject’s speech was described as “incoherent”, “disorganized”, or “flight-of-ideas”. Cases of Charles-Bonnet syndrome, or visual-release hallucinations that occur as a complication of acquired visual field defects and occur exclusively in the blind field of vision, were excluded because these types of hallucinations are not seen in schizophrenia.

*Relationship to Previous Lesion Network Mapping Studies*

Lesion Network Mapping (LNM) has previously been used to establish a model of familiarity detection by mapping the functional connectivity of lesions that cause isolated delusions of misidentification^3^. This study successfully identified regions specific to familiarity but specifically excluded other types of delusions, specifically persecutory delusions, that are more common in schizophrenia. A second study identified a common circuit for lesions that cause hallucinations across sensory modalities (visual, auditory, olfactory, tactile)^4^. Olfactory and tactile hallucinations are rare in schizophrenia, and most (78/89) cases from this study presented with a single psychotic symptom and therefore would not meet criteria for schizophrenia. Thus, while both studies investigated important anatomy for isolated symptoms and general hallucinations, neither attempted to model schizophrenia.

**Lesion Network Mapping**

*Lesion Reconstruction*

Lesions were reconstructed by manually tracing the published images onto a common MNI template using 3D Slicer (eFigure 2)(Boston, MA; https://www.slicer.org/)^5^. Although we did not have access to the original image files and published cases provide only 2-dimensional images of the lesions, prior work has demonstrated that this approximation is sufficient for LNM studies^3^.

*Determination of Functional Connectivity of Lesion Locations*

The functional connectivity of each lesion location was estimated using a normative database of resting-state functional connectivity (n=1000) ^6^, generating a brain-wide map of correlation values for each lesion. We derived maps of brain-wide t-values for each lesion by performing a one-sample t-test on the Fisher-transformed correlation values derived from the correlation values between the lesion location and the database of resting-state functional connectivity. These analyses were repeated and validated using a comparable normative database of resting-state functional connectivity that is publicly available (sensitivity analysis r= 0.97, specificity analysis r= 0.98. Both analyses identified the same peak regions regardless of which connectome was used)(Brain Genomics Superstruct Project, <https://dataverse.harvard.edu/dataverse/GSP>) ^7^. The normative connectome data were preprocessed as previously described^6,7^.
The connectome employed in this study is derived from healthy individuals, not from individuals with psychotic symptoms. Studying the connectivity of lesions using a connectome derived from individuals with psychosis could yield results of interest. However, subjects included in this study had no history of psychosis and developed psychosis acutely following a lesion; therefore, a connectome derived from healthy subjects would be a better approximation of the connectivity of the subjects in this study and allow for the most direct interpretation of our findings.

*Sensitivity Analysis*

Functional connectivity maps were thresholded at |t|>7 to remain consistent with the methodology of prior lesion-network mapping studies^8^. However, these studies have sometimes been criticized for arbitrary thresholding^9,10^. This method is also limited by the fact that it provides a metric of overlap percentage but does not account for variance between affected lesions. To improve upon this established method, we conducted an additional layer of unthresholded testing by using a one-sample t-test on the functional connectivity maps. This not only eliminates concern about the arbitrary threshold, but also enables voxel-wise statistical testing that takes variance into account. To be consistent with prior literature and avoid arbitrary thresholding, we only included voxels that passed both tests.

*Specificity Analysis*

We evaluated the specificity of our findings by comparing the connectivity maps of lesions that caused psychosis with two independent control groups: lesions of consecutive stroke patients presenting to Massachusetts General Hospital (n=490)^11^, and lesions in our group’s database that caused symptoms not related to psychosis (n=666, including consecutive incidental strokes, Addiction, Akinetic Mutism, Alien Limb syndrome, Amnesia, Anton syndrome, Asterixis, Blindsight, Coma, Confabulations, Cortical blindness, Capgras Delusions, Depression, Dystonia, Freezing of Gait, Hemichorea, Holmes Tremor, Infantile Spasms, Loss of Consciousness, Pain, Parkinsonism, Prosopagnosia, and Vertigo). Because psychosis is a rare complication of stroke^12^ we anticipate the majority of these patients did not have psychotic symptoms. While there may be some patients with unreported psychotic symptoms in our control group, this would bias our analyses against finding a positive result. Thus, control lesions with unreported psychotic symptoms are unlikely to artificially inflate our reported results.

We performed two-sample t-tests at each voxel between the cases and combined controls group (n=1156). We used the permutation-based FWE correction in FSL PALM^13^ to correct for multiple comparisons, following the Westfall-Young procedure. The voxels most specific to lesions that cause psychosis were in the posterior subiculum of the hippocampus. This result was driven by the experimental group rather than the control group, as the peak voxel in the subiculum (MNI [24,-24,-16], T=15.5) from this two-sample t-test was not strongly represented in one-sample t-tests of the control group (T=0.6). Results were highly similar when comparing lesions that caused psychosis with each set of controls separately (r=0.90)(eFigure 4).

*Conjunction*

We binarized the sensitivity map according to whether a voxel was connected to at least 75% of lesions causing psychosis at a threshold of |T| ≥ 7 and also survived the FWE-corrected one-sample t-test. We binarized the results of the specificity analysis according to whether a voxel survived FWE correction. We overlaid these binarized maps and considered voxels that were commonly nonzero in both maps to be sensitive and specific.

We identified several secondary peak regions at slightly lower thresholds for sensitivity (overlap >75%) and specificity (p_FWE_<5x10^-4^) in the psychosis circuit: the mediodorsal and midline nuclei of the thalamus, the VTA, the retrosplenial cortex, and lobule IX and the dentate nucleus of the cerebellum (eFigure 5). Outside the cerebellar findings, all of these regions have robust white matter connections with the hippocampus^14-22^. Lobule IX and the dentate nucleus of the cerebellum are not directly connected to the hippocampus. Lobule IX has been associated with language, social, and emotional task performance^23^ and is a principal region of cerebellar grey matter loss in patients with schizophrenia^24^. The dentate nucleus has been implicated in previous LNM studies of auditory hallucinations^4^. It is notably the only cerebellar output known to affect the prefrontal cortex, and does so in part through projections to the mediodorsal nucleus of the thalamus^25^.

Interestingly, recent work has identified a significant correlation between retrosplenial cortex-specific gene expression and schizophrenia, providing insight into the cellular profiles related to this diagnosis^26^.

The involvement of the VTA in this psychosis circuit is consistent with the established dopamine hypothesis of schizophrenia — i.e dopaminergic agonists can cause psychosis, while dopaminergic antagonists are the first-line treatment for schizophrenia. Neurotransmitters such as dopamine have widespread connections throughout the brain and could account for the coordination of disparate brain regions^2^^7^. In addition to the VTA, several regions in this psychosis circuit are involved in dopaminergic communication. Notably, the subiculum’s capacity to modulate dopaminergic neurons in the VTA^2^^8^ is hypothesized to explain pathologically increased salience in patients with schizophrenia^2^^9^. Further, the mediodorsal nucleus highly expresses dopamine receptors^30,31^, and the dentate nucleus has a role in regulating dopaminergic neurotransmission^32^.

**Analysis of Covariates**

To account for age and sex, we performed a partial correlation including these covariates. However, these data were not systematically collected in most of the control cohorts. We employed the two subsets of control cohorts that had age and sex covariates available (lesions that caused infantile spasms (n = 74)^33^ and a cohort of incidental stroke lesions that caused various symptoms but did not cause psychosis (n = 135)^34^. We then performed a voxel-wise partial correlation analysis accounting for age and sex. The functional connectivity topography of the resulting map had a very high spatial correlation with the map generated from all of the control groups without incorporating age and sex covariates (r = 0.98), and peak voxels remained in the subiculum (eFigure 6).

**Lesion Network Mapping without Hippocampal Lesions**

We repeated the above analysis after removing any lesions that had voxels in the hippocampus as defined by the Brainnetome atlas^35^. In the cohort of lesions that did not touch the hippocampus, we then repeated the LNM analyses as described above. For the sensitivity test, we calculated which voxels had overlapping functional connections of >75% of non-hippocampal lesions at a threshold of |T| ≥ 7. We then overlaid an FWE-corrected one-sample t-test of the functional maps of non-hippocampal lesions, and only included voxels that passed both tests in our sensitivity result. For the specificity test, we compared the functional maps of non-hippocampal lesions that caused psychosis with the same control group described above (eFigure 7). In addition, we examined how many lesions included a voxel within the most sensitive and specific region of the psychosis circuit (posterior subicular region, shown in Figure 1), and found 20 cases (13%) had at least one voxel in this region.

**Similarity Between Different Symptoms of Psychosis**

We first categorized cases into five classes of positive symptoms of psychosis: delusions, thought disorder, visual hallucinations, auditory hallucinations, and negative symptoms. If a case had more than one symptom, they were included in multiple symptom groups.

We derived specificity maps by performing a voxel-wise two-sample t-test between the functional connectivity of lesion locations of symptom group and all the lesions that were not part of their symptom group. This was done for each symptom of psychosis and each subgroup within the control group (Consecutive incidental strokes, Addiction, Akinetic Mutism, Alien Limb syndrome, Amnesia, Anton syndrome, Asterixis, Blindsight, Coma, Confabulations, Cortical blindness, Criminality, Capgras Delusions, Depression, Dystonia, Epilepsy, Freezing of Gait, Hemichorea, Holmes Tremor, Infantile Spasms, Loss of Consciousness, Mania, Pain, Parkinsonism, Prosopagnosia, and Vertigo). We then performed spatial correlations between each of these specificity maps and the general psychosis circuit. If we compared a psychotic symptom group and the general psychosis circuit, we excluded lesions from the psychotic symptom group being examined. We then performed a LNM analysis on each of the psychotic symptom groups, with thresholds of >75% overlap of functional connections for the sensitivity test and p_FWE_<5 x 10^-4^ for the voxel-wise two-sample t-test against controls for the specificity test.

To determine the similarity between the functional connectivity profiles of each individual lesion and the psychosis circuit, we performed a leave-one-out cross-validation. A spatial correlation was computed between the functional connectivity profile of each individual lesion (eFigure 8a) and a specificity map generated from the remaining 152 lesions (eFigure 8b). The overall functional connectivity profile was similar between lesions that cause psychotic symptoms (eFigure 8c). We tested for significance by randomly permuting case and control labels to produce 10,000 two-sample t-tests. We then computed the distribution of spatial correlations between the functional connectivity maps of each case lesion and the maps from the randomly permuted two-sample t-test. The p-value was then computed as the proportion of permuted means that were greater than or equal to the observed mean (see Non-Parametric Results, below).

To further test the hypothesis that lesions that caused different symptoms of psychosis were more similar to each other than to controls, we used spatial correlations to compare the functional maps of each symptom group (Figure 3a) to a one-sample t-test of control lesions and a one-sample t-test of psychosis lesions, excluding the symptom under examination. (Figure 3b). We performed a two-sample t-test between the distributions of correlations with controls and the distributions of correlations with other lesions that cause psychosis (Figure 3c). The functional connectivity of lesions that caused each psychotic symptom was more similar to each other to lesions that did not cause psychosis (Figure 3d). We additionally performed a non-parametric test by comparing the distribution of correlation values between the functional maps of each symptom group and the one-sample t-test of the other psychotic symptoms, and the distribution of correlation values between the functional maps of each symptom group and the  one-sample t-test of the randomly selecting an controls. The number of controls that were randomly selected were proportional to the comparative group for each symptom cohort (see Non-Parametric Results, below).

From this analysis we conclude that different psychotic symptoms converge on a common network. The posterior subiculum was not specific to any particular symptom of psychosis but may instead be a common functional connection of schizophrenia-like symptoms.

**Distinctions between Different Symptoms of Psychosis Presenting as a Psychotic Syndrome**

In an exploratory analysis, we compared each symptom-specific grouping with the remaining psychosis-inducing lesions using a voxel-wise partial correlation, controlling for co-occurring symptoms in each case. We used the permutation-based correction available in FSL PALM for multiple comparisons (p_FWE_ < 0.05). In most cases, the functional maps of lesions that caused each group of psychotic symptoms did not significantly differ from the functional maps of lesions that caused other psychotic symptoms. The exception was lesions that caused thought disorder (n=25), which were significantly more positively correlated to the fornix, body of the corpus callosum, and a cluster of voxels extending from the anterior corona radiata to the inferior frontal gyrus bilaterally (eFigure 9). Notably, lesions that caused symptoms described as “schizophrenia” or “schizophrenia-like” did not significantly differ from other cases of lesions that caused psychotic symptoms, indicating that these two groupings shared similar functional connectivity profiles. These results suggest that lesions causing multiple psychotic symptoms have a common functional connectivity profile.

**Distinctions between Different Symptoms of Psychosis in Isolation**

To determine if isolated psychotic symptoms and the same symptoms presenting as part of a syndrome mapped to distinct brain regions, we examined each cohort of lesions that caused psychotic symptoms in isolation. We performed LNM on each of these cohorts, with thresholds of >85% overlap of functional connections at a threshold |T|>7 for the sensitivity test and p_FWE_<0.05 for the voxel-wise two-sample t-test against controls for the specificity test. In addition to FWE correction for comparisons across voxels, we used a False Discovery Rate correction to account for comparative analysis across the symptom groupings. As discussed above, cases of Charles Bonnet syndrome were excluded from this study. Among cases of isolated auditory hallucinations, six had hallucinations of simple repetitive sounds (e.g. “rain falling on the roof” or “a machine humming”), 14 heard familiar music, and four had both familiar music and simple sounds. None of these patients received a psychiatric diagnosis, and none were included in the schizophrenia(-like) cohort in this study. All 8 cases of isolated delusions had paranoid delusions. Isolated visual hallucinations mapped to the subiculum of the hippocampus, isolated auditory hallucinations mapped cortically to the superior temporal gyrus and subcortically to the subcortical auditory system, and isolated delusions mapped to the mediodorsal nucleus of the thalamus. Corroborating the mediodorsal nucleus findings, this region of the thalamus has previously been implicated in causal studies using animal models of paranoid delusions^36,37^.

**Distinguishing Circuits of Lesions that Cause Amnesia and Lesions that Cause Amnesia with Psychosis**

The anatomical similarity of the psychosis and memory circuits brings into question why apparently similar lesions can cause psychosis in some patients but amnesia in others. Memory deficits are a known complication of schizophrenia^38,39^, but psychosis does not always accompany memory deficits. We first examined the 53 cases used to derive a memory circuit^40^ to identify if any of them had psychotic symptoms. We also examined the 153 cases used for the psychosis circuit to determine how many had amnestic symptoms. Among the 53 cases of lesions causing amnesia, six were excluded for having co-occurring psychotic symptoms (one had visual hallucinations, one had auditory hallucinations, one had delusions, two had thought disorder, one had visual hallucinations and delusions). Among the 153 cases of lesion-induced psychosis in this study, 48 reported an acute memory deficit, 7 reported no memory deficit, and 98 did not report any memory testing.

We then compared the functional connectivity of lesions that caused psychosis and amnesia with the functional connectivity of lesions that just caused amnesia using a two-sample t-test at each voxel. We used the FWE method (p_FWE_ < 0.05) to correct for multiple comparisons.

To determine if the laterality of lesions to different cerebral hemispheres could distinguish between lesions that cause psychosis and amnesia and lesions that cause just amnesia, we used the Brainnetome Atlas to determine the number of voxels each lesion had in each hemisphere. We then categorized lesions based on whether they were located in the right hemisphere, the left hemisphere, or both and performed a 𝝌^2^ test to determine significance.

Comparing lesions that cause psychosis and amnesia with lesions that cause just amnesia, lesions causing psychosis were more functionally connected to aspects of the superior temporal gyrus, the ventral claustrum, and part of the uncinate fasciculus (p_FWE_ < 0.05) (eFigure 10). Lesions causing amnesia without psychosis were not associated with any significant voxels in this analysis. Comparing all lesions that cause psychosis with lesions that cause just amnesia, lesions causing psychosis were more functionally connected to aspects of the middle temporal gyrus, and lesions causing amnesia were more functionally connected to the retrosplenial cortex, anterior nuclei of the thalamus, fornix, and medial prefrontal cortex (p_FWE_ < 0.05).

Lesions that cause psychosis were more likely to have functional connections to the temporal gyri, though more detailed symptoms for each case would likely improve anatomical distinction. Lesions that cause psychosis were also more likely to be restricted to the right hemisphere, consistent with literature reporting the development of psychotic symptoms from right-sided lesions in stroke patients^12^ and right-sided hippocampal damage in patients with traumatic brain injury^41^ and Alzheimer’s disease^42^.

**Validation in Independent Cohort**

We used the Vietnam Head Injury Study (VHIS)^43^ for external validation. This dataset has lesions from 181 Veterans (mean age 58.4 ± 3.1) who completed the neurobehavioral rating scale subsequent to penetrating brain trauma during the Vietnam War. Of the categories in the behavioral rating, “suspiciousness”, “unusual thought content”, “hallucinatory behavior”, and “conceptual disorganization” describe positive symptoms of psychosis. “Hallucinatory behavior” was excluded from our analysis because only 2 of 181 patients in the VHIS were rated as displaying hallucinatory behavior, and both were given a 1 (“very mild”) out of 6 on a Likert scale. “Conceptual disorganization” was included in the analysis but was less specific than “unusual thought content” as a psychotic symptom because all the participants who were rated as having “unusual thought content” (n=21) were also rated as having “conceptual disorganization”, but many patients rated as having “conceptual disorganization” were not rated with “unusual thought content” (n=41/62). The criteria for “conceptual disorganization” included symptoms not specific to psychosis such as tangential and perseverative speech. We therefore considered “unusual thought content” and “suspiciousness” as psychosis-like symptoms, and treated the other categories as controls. We estimated the functional connectivity of each lesion using the same normative database of resting-state functional connectivity (n=1000) as in our previous analyses. We correlated the functional maps of these lesions with their associated behavioral ratings and the map produced from the LNM specificity analysis, yielding a correlation between the psychosis circuit and the behavioral rating associated with brain lesions in an independent dataset. We hypothesized that lesions with greater connectivity to the psychosis circuit would be associated with psychosis-like symptoms but not other symptoms. Additionally, we employed non-parametric testing using 10,000 permutations to generate a null model. For each permutations, patients' imaging were randomly shuffled and the correlation of each behavioral rating and the psychosis specificity map was recalculated. P-values were computed by comparing the actual correlations with the distribution of 10,000 permutations (see Non-Parametric Results, below).

To investigate if the ‘suspiciousness’ and ‘unusual thought content’ functional connectivity maps converged on a similar regions as the psychosis circuit (i.e. the posterior subiculum), we first performed a voxel-wise regression analysis using the functional connectivity map and the Neurobehavioral Rating Scale score for ‘suspiciousness’ and ‘unusual thought content’ corresponding to each patient. This resulted in a correlation coefficient map representing the association between voxel-wise connectivity and patient scores. While the overall ‘suspiciousness’ and ‘unusual thought content’ circuits were significantly associated with the psychosis circuit compared to controls, no individual region in either map survived multiple comparisons correction using permutation-based FWE correction. This could be due to noise in the VHIS dataset, or because patients hippocampal lesions would be more likely to have cognitive impairment and thus be more likely to lack capacity to consent to participation in the VHIS study.

**Non-parametric Results:**

Leave-one-functional-map-out analysis (eFigure 8): (p<0.0001).

Similarity Between Different Symptoms of Psychosis (eFigure 3): Similar to the parametric testing, the functional connectivity of lesions that caused each psychotic symptom were more similar to each other than to lesions that did not cause psychosis (auditory hallucinations: p<0.0001, visual hallucinations: p=0.0006, thought disorders: p=0.0101, negative symptoms: p=0.0003 described as “schizophrenia”: p<0.0001).

Validation in Independent Cohort (Figure 4):Of the 23 behavioral ratings, ‘suspicious’ and ‘unusual thought content’ remained the only two significant behavioral ratings (p=0.0091 and p=0.0185, respectively).

**Lesion Network Mapping - Determination of Therapeutic Target**

To determine a therapeutic target, we constructed a precomputed functional connectome by computing average Blood Oxygenation Level Dependent (BOLD) signal correlations for each voxel across 1,000 participants. This generated a brain-wide functional map distinct for each voxel showing the BOLD correlations with the other 228,453 voxels in a 2mm-resolution brain atlas. We then correlated the functional map of each voxel with the functional map generated by the specificity test to look for the voxels whose BOLD correlations across the brain most closely resembled the functional map specific to lesions that cause psychosis. This method has retrospectively replicated known therapeutic neuromodulation targets for tics^44^, depression^45^ and addiction^46^, but has not yet been used prospectively to identify novel therapeutic targets. Notably, this method is intended to target a brain-wide circuit and ameliorate behavioral symptoms. There is extensive research into using functional correlations to derive surface TMS targets to modulate deep structures such as the hippocampus^47,48^. Future studies could compare these methods to determine differential effects.

**Sham-Controlled TMS Trials for Schizophrenia:**

Multiple sham-controlled TMS trials for schizophrenia have been conducted, predominantly targeting the DLPFC, cerebellar vermis, or temporoparietal junction. The DLPFC and cerebellar targets have shown improvements in negative, but not positive, symptoms in the active group compared to sham^49-53^. Some TMS studies targeting the temporoparietal junction have demonstrated reduction in auditory hallucinations^54-63^, but this finding has not been consistently replicated^64-69^.

**Hippocampus and Psychosis Literature**

The anterior CA1 subfield of the hippocampus has been shown to have increased blood volume – a marker of hyperactivity - in patients with a psychosis-spectrum disorder^70-72^ , and this abnormality may progress to the subiculum when clinical high-risk patients transition to psychosis^73^. Interestingly, increased blood volume in anterior CA1 is associated with impaired activity of anterior hippocampal neurons during memory tasks^74.^ Some post-mortem studies suggest neurotransmitter markers of hyperactivity in CA3 that then progress to CA1^75^. The anterior and posterior aspects of the hippocampus can be functionally distinguished, as the anterior aspect is preferentially engaged in associational memory and the posterior aspect more engaged in spatial/contextual memory^76-78^. Schizophrenia has been associated with abnormal anterior/posterior hippocampal recruitment on task-based fMRI^74,79^, and early-stage psychotic illness is associated with abnormally increased functional connectivity within the hippocampal network^80,81^.

### eTables

**eTable 1**

| **Author and date** | **Patient number within study** | **Sex** | **Age** | **Visual Hallucination** | **Auditory Hallucination** | **Delusion** | **Thought Disorder** | **Described as Schizophrenia or Schizophrenia-like** | **Memory Deficits** | **Multiple Symptoms** |
| --- | --- | --- | --- | --- | --- | --- | --- | --- | --- | --- |
| **Acioly 2010** |  | W | 32 |  |  |  |  |  | Impaired Memory | ≡ |
| **Allan Hobson 2002** |  | M | 67 |  |  |  |  |  | Not Described | • |
| **Almeida 2011** |  | W | 12 |  |  |  |  |  | Not Described | ≡ |
| **Arasappa 2013** |  | M | 38 |  |  |  |  |  | Not Described | ≡ |
| **Arikan 2009** |  | M | 33 |  |  |  |  |  | No Deficits | ≡ |
| **Asghar-Ali 2004** | Case 1 | W | 54 |  |  |  |  |  | Impaired Memory | ≡ |
|  | Case 2 | W | 50 |  |  |  |  |  | Impaired Memory | ≡ |
| **Barboza 2013** |  | W | 54 |  |  |  |  |  | Not Described | ≡ |
| **Benke 2006** | Case 2 | M | 46 |  |  |  |  |  | Not Described | ≡ |
|  | Case 3 | M | 41 |  |  |  |  |  | Impaired Memory | ≡ |
|  | Case 4 | W | 74 |  |  |  |  |  | Impaired Memory | ≡ |
| **Bielawski 2015** |  | M | 56 |  |  |  |  |  | Impaired Memory | ≡ |
| **Breitner 1990** | Case 1 | W | 75 |  |  |  |  |  | Impaired Memory | ≡ |
|  | Case 8 | W | 86 |  |  |  |  |  | Not Described | ≡ |
| **Calabrò 2012** |  | W | 82 |  |  |  |  |  | No Deficits | • |
| **Carmona-Bayonas 2017** |  | W | 60 |  |  |  |  |  | Not Described | ≡ |
| **Carson 1997** |  | M | 9 |  |  |  |  |  | Not Described | ≡ |
| **Cascino 1986** | Case 1 | W | 32 |  |  |  |  |  | No Deficits | • |
|  | Case 2 | W | 42 |  |  |  |  |  | Not Described | • |
|  | Case 3 | W | 53 |  |  |  |  |  | Not Described | • |
| **Castaño Ramírez 2020** |  | W | 60 |  |  |  |  |  | Impaired Memory | ≡ |
| **Cerrato 2001** |  | M | 35 |  |  |  |  |  | Not Described | • |
| **Chaudhari 2018** |  | M | 16 |  |  |  |  |  | Not Described | ≡ |
| **Chrispal 2009** |  | W | 72 |  |  |  |  |  | Not Described | • |
| **Cohen 1992** |  | M | 30 |  |  |  |  |  | Impaired Memory | • |
| **Cosentino 2010** |  | M | 63 |  |  |  |  |  | Not Described | • |
| **de la Fuente Fernandez 1994** |  | W | 70 |  |  |  |  |  | No Deficits | • |
| **Dinges 2013** | Case 1 | W | 89 |  |  |  |  |  | Not Described | • |
|  | Case 2 | W | 75 |  |  |  |  |  | Not Described | • |
| **Dobry 2014** |  | M | 51 |  |  |  |  |  | Impaired Memory | ≡ |
| **Dogan 2013** |  | W | 63 |  |  |  |  |  | Not Described | ≡ |
| **Douen 1997** |  | M | 43 |  |  |  |  |  | Not Described | • |
| **Elkhaled 2020** |  | M | 23 |  |  |  |  |  | Not Described | ≡ |
| **Estronza 2018** |  | M | 42 |  |  |  |  |  | Not Described | ≡ |
| **Feinberg 1989** |  | M | 83 |  |  |  |  |  | Not Described | ≡ |
| **ffytche 2003** |  | W | 78 |  |  |  |  |  | Not Described | • |
| **Gabelić 2012** |  | M | 25 |  |  |  |  |  | Not Described | ≡ |
| **Geddes 2016** |  | M | 66 |  |  |  |  |  | No Deficits | • |
| **Geller 1987** |  | M | 61 |  |  |  |  |  | Not Described | • |
| **Ghosh 2015** |  | W | 71 |  |  |  |  |  | No Deficits | • |
| **Gilberthorpe 2017** | Case 7 | M | 50 |  |  |  |  |  | Impaired Memory | ≡ |
| **Godani 2012** |  | W | 67 |  |  |  |  |  | Not Described | • |
| **Guptha 2007** |  | M | 62 |  |  |  |  |  | Not Described | ≡ |
| **Haas 2019** |  | W | 62 |  |  |  |  |  | Not Described | ≡ |
| **Haddad - 2021** |  | W | 53 |  |  |  |  |  | Impaired Memory | ≡ |
| **Hall 1992** |  | M | 23 |  |  |  |  |  | Impaired Memory | ≡ |
| **Harris_1991** |  | M | 65 |  |  |  |  |  | Not Described | • |
| **Hatano 2019** |  | M | 65 |  |  |  |  |  | Not Described | ≡ |
| **Hattori 1988** | Case 1 | M | 73 |  |  |  |  |  | Not Described | • |
| **Howlett 1994** |  | M | 63 |  |  |  |  |  | Not Described | • |
| **Ido 2018** |  | W | 56 |  |  |  |  |  | Not Described | ≡ |
| **Ishii 2011** |  | M | 21 |  |  |  |  |  | Not Described | ≡ |
| **Isolan 2010** |  | M | 34 |  |  |  |  |  | Not Described | • |
| **Jiang 2018** |  | M | 23 |  |  |  |  |  | Not Described | ≡ |
| **Julayanont 2017** |  | M | 55 |  |  |  |  |  | Not Described | ≡ |
| **Kadak 2013** |  | M | 13 |  |  |  |  |  | Impaired Memory | ≡ |
| **Kamalakannan 2004** |  | M | 84 |  |  |  |  |  | Not Described | • |
| **Kan 1989** |  | F | 38 |  |  |  |  |  | Not Described | ≡ |
| **Karakula-Juchnowicz 2018** |  | W | 27 |  |  |  |  |  | Not Described | ≡ |
| **Kasabkojian 2021** |  | M | 55 |  |  |  |  |  | Impaired Memory | ≡ |
| **Keshavan 1988** |  | M | 43 |  |  |  |  |  | Not Described | ≡ |
| **Kho 2007** |  | W | 23 |  |  |  |  |  | Not Described | ≡ |
| **Khong 2007** |  | W | - |  |  |  |  |  | Impaired Memory | ≡ |
| **Kim 2015** |  | M | 45 |  |  |  |  |  | Impaired Memory | • |
| **Klasen 1999** |  | M | 12 |  |  |  |  |  | Impaired Memory | ≡ |
| **Kobayashi 2018** |  | W | 67 |  |  |  |  |  | Impaired Memory | • |
| **Kolmel_1991** |  | M | 56 |  |  |  |  |  | Not Described | ≡ |
| **Kosman 2018** |  | W | 52 |  |  |  |  |  | Impaired Memory | ≡ |
| **Kumral 2001** |  | W | 75 |  |  |  |  |  | Impaired Memory | • |
| **Kumral 2015** | Case 4 | - | - |  |  |  |  |  | Impaired Memory | • |
|  | Case 5 | - | - |  |  |  |  |  | Impaired Memory | • |
|  | Case 8 | - | - |  |  |  |  |  | Impaired Memory | • |
|  | Case 9 | - | - |  |  |  |  |  | Impaired Memory | • |
|  | Case 12 | - | - |  |  |  |  |  | Impaired Memory | • |
|  | Case 13 | - | - |  |  |  |  |  | Impaired Memory | • |
|  | Case 14 | - | - |  |  |  |  |  | Impaired Memory | • |
| **Lanska 1987** |  | M | 55 |  |  |  |  |  | Not Described | • |
| **Lee 2011** |  | M | 20 |  |  |  |  |  | Impaired Memory | • |
| **Leeks 1967** |  | M | 28 |  |  |  |  |  | Impaired Memory | ≡ |
| **Leo 2016** |  | W | 59 |  |  |  |  |  | Impaired Memory | ≡ |
| **Li 2018** |  | W | 44 |  |  |  |  |  | Not Described | • |
| **Liao 2018** |  | W | 67 |  |  |  |  |  | Not Described | • |
| **Lim 2011** |  | M | 44 |  |  |  |  |  | Not Described | • |
| **Lisanby 1998** |  | W | 26 |  |  |  |  |  | Not Described | ≡ |
| **LoBuono 2019** | Case 1 | W | 55 |  |  |  |  |  | Impaired Memory | ≡ |
|  | Case 2 | W | 45 |  |  |  |  |  | Impaired Memory | ≡ |
| **Luauté 2008** |  | M | 77 |  |  |  |  |  | Impaired Memory | ≡ |
| **Maiuri 2002** | Case 1 | W | 69 |  |  |  |  |  | Not Described | • |
|  | Case 2 | W | 44 |  |  |  |  |  | Not Described | • |
| **Mäkelä 1997** | Case 5 | M | 59 |  |  |  |  |  | Impaired Memory | • |
|  | Case 6 | M | 54 |  |  |  |  |  | Impaired Memory | • |
| **Malamud 1967** | Case 13 | W | 43 |  |  |  |  |  | Impaired Memory | • |
| **McGilchrist 1993** |  | M | 43 |  |  |  |  |  | Impaired Memory | ≡ |
| **McKee 1990** |  | M | 83 |  |  |  |  |  | Impaired Memory | ≡ |
| **McMurtray 2014** | Case 1 | M | 59 |  |  |  |  |  | Not Described | ≡ |
|  | Case 2 | W | 52 |  |  |  |  |  | Not Described | ≡ |
| **Minabe 1990** |  | F | 40 |  |  |  |  |  | Not Described | ≡ |
| **Mittal 2010** |  | W | 19 |  |  |  |  |  | Not Described | ≡ |
| **Mittal 2010** |  | M | 17 |  |  |  |  |  | Not Described | ≡ |
| **Miyazawa 2001** |  | W | 53 |  |  |  |  |  | Not Described | • |
| **Miyazawa 2009** |  | W | 68 |  |  |  |  |  | Not Described | • |
| **Mizukami 1999** |  | M | 18 |  |  |  |  |  | Impaired Memory | ≡ |
| **Mocellin 2006** | Case A | W | 85 |  |  |  |  |  | No Deficits | • |
|  | Case B | W | 68 |  |  |  |  |  | Not Described | ≡ |
| **Mollet 2007** |  | W | 61 |  |  |  |  |  | Impaired Memory | ≡ |
| **Morris 2013** |  | W | 22 |  |  |  |  |  | Not Described | ≡ |
| **Murata 1994** |  | M | 55 |  |  |  |  |  | Not Described | • |
| **Nagaratnam 1996** |  | W | 84 |  |  |  |  |  | Not Described | ≡ |
| **Narumoto 2005** |  | W | 55 |  |  |  |  |  | Impaired Memory | ≡ |
| **Nishio 2007** |  | W | 74 |  |  |  |  |  | Not Described | ≡ |
| **Noda 1993** | Case 1 | M | 72 |  |  |  |  |  | Not Described | ≡ |
|  | Case 2 | W | 46 |  |  |  |  |  | Impaired Memory | ≡ |
| **Notas 2015** |  | M | 79 |  |  |  |  |  | Not Described | • |
| **Oladiran 2018** |  | M | 28 |  |  |  |  |  | Not Described | • |
| **Pant 1996** |  | M | 22 |  |  |  |  |  | Not Described | ≡ |
| **Paquier 1992** |  | W | 52 |  |  |  |  |  | Not Described | • |
| **Pavesi 2014** |  | W | 48 |  |  |  |  |  | Not Described | • |
| **Preuss 2009** |  | F | 60 |  |  |  |  |  | Not Described | • |
| **Reiss 2006** |  | W | 35 |  |  |  |  |  | Not Described | ≡ |
| **Roberts 1990** | Case 60 | - | - |  |  |  |  |  | Not Described | ≡ |
|  | Case 99 | - | - |  |  |  |  |  | Not Described | ≡ |
|  | Case 102 | - | - |  |  |  |  |  | Not Described | ≡ |
|  | Case 144 | - | - |  |  |  |  |  | Not Described | ≡ |
| **Roberts 2001** |  | W | 61 |  |  |  |  |  | Not Described | • |
| **Rousseaux 1994** |  | M | 15 |  |  |  |  |  | Not Described | • |
| **Russel 2003** | Case 1 | W | 28 |  |  |  |  |  | Not Described | • |
|  | Case 2 | W | 27 |  |  |  |  |  | Not Described | • |
| **Sade 2006** |  | W | 38 |  |  |  |  |  | Not Described | ≡ |
| **Sato 1993** |  | W | 55 |  |  |  |  |  | Not Described | ≡ |
| **Schielke 2000** |  | M | 57 |  |  |  |  |  | Not Described | • |
| **Schwartz 2013** |  | W | 28 |  |  |  |  |  | Impaired Memory | ≡ |
| **Serby 2013** | Case 1 | W | 79 |  |  |  |  |  | Not Described | • |
|  | Case 3 | W | 85 |  |  |  |  |  | Not Described | • |
|  | Case 2 | M | 84 |  |  |  |  |  | Not Described | • |
| **Serra Catafau 1992** |  | M | 68 |  |  |  |  |  | Not Described | • |
| **Singh 2011** |  | W | 15 |  |  |  |  |  | Not Described | ≡ |
| **Struck 1992** |  | W | 19 |  |  |  |  |  | Not Described | ≡ |
| **Suradom 2020** |  | W | 51 |  |  |  |  |  | Not Described | ≡ |
| **Suzuki 2003** |  | M | 58 |  |  |  |  |  | Impaired Memory | ≡ |
| **Talih 2013** |  | M | 70 |  |  |  |  |  | Not Described | • |
| **Tanriover 2008** |  | W | 25 |  |  |  |  |  | Not Described | • |
| **Taylor 2005** |  | W | 24 |  |  |  |  |  | Not Described | ≡ |
| **Thomas 2017** |  | M | 61 |  |  |  |  |  | Impaired Memory | ≡ |
| **Udaya 2015** |  | W | 21 |  |  |  |  |  | Not Described | ≡ |
| **Vita 2008** |  | M | 11 |  |  |  |  |  | Not Described | ≡ |
| **Wong 1993** |  | W | 30 |  |  |  |  |  | Not Described | ≡ |
| **Wong 2011** |  | W | 45 |  |  |  |  |  | Not Described | ≡ |
| **Woo 2014** |  | W | 85 |  |  |  |  |  | Impaired Memory | • |
| **Yadav 2010** |  | M | 18 |  |  |  |  |  | Not Described | ≡ |
| **Yalcin 1996** | Case 1 | W | 48 |  |  |  |  |  | Not Described | ≡ |
|  | Case 2 | M | 51 |  |  |  |  |  | Impaired Memory | • |
|  | Case 3 | M | 51 |  |  |  |  |  | Not Described | • |
| **Zhou 2020** |  | W | 76 |  |  |  |  |  | Not Described | ≡ |

≡ Multiple psychotic symptoms

• Single psychotic symptom

### eFigures

**eFigure 1**

Results of PRISMA-formatted literature review

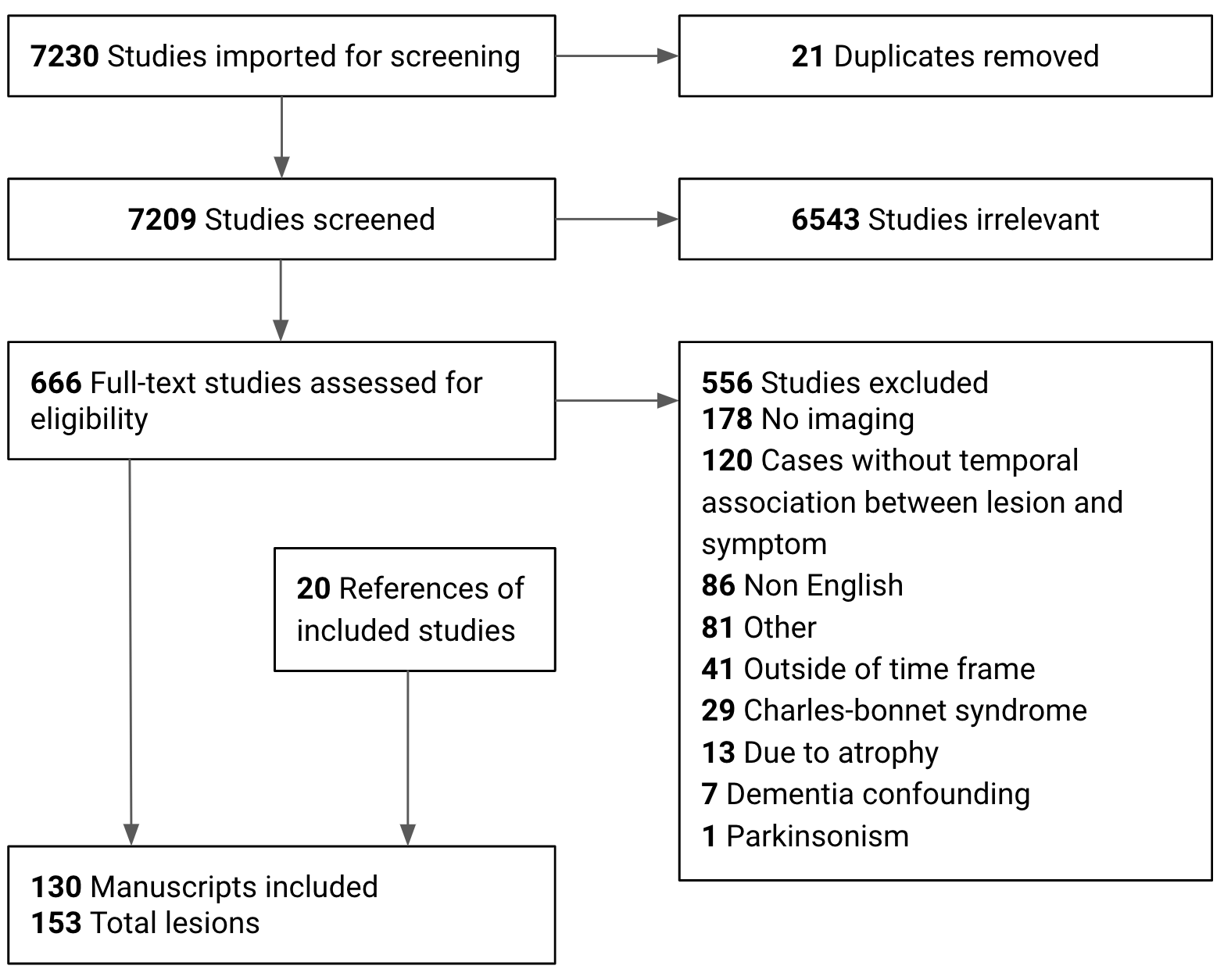

We identified lesions causing psychosis via a PubMed search including case reports, reviews, and systematic reviews published June, 1957- March, 2021 using the following search terms for All Fields:

(psychosis OR Schizophrenia OR hallucination OR hallucinosis) AND (lesion OR tumor OR tumour OR stroke OR infarct OR ischemic OR hemorrhage OR haemorrhage OR bleeding OR traumatic)

**eFigure 2**

Example Lesions Associated with Psychosis (6 of 153)

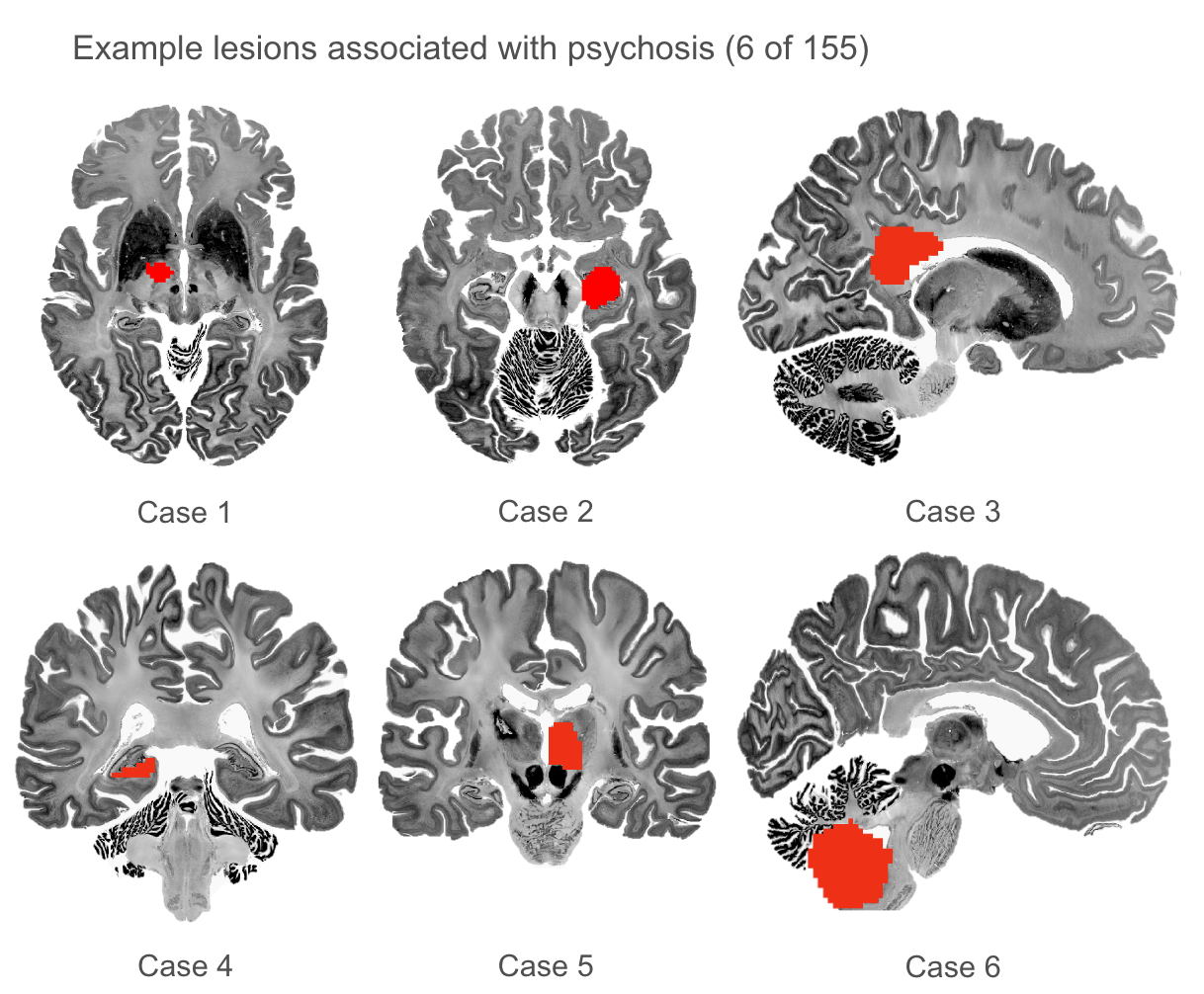

**eFigure 3**

Lesions associated with psychosis and control lesions were distributed throughout vascular territories and brain regions.

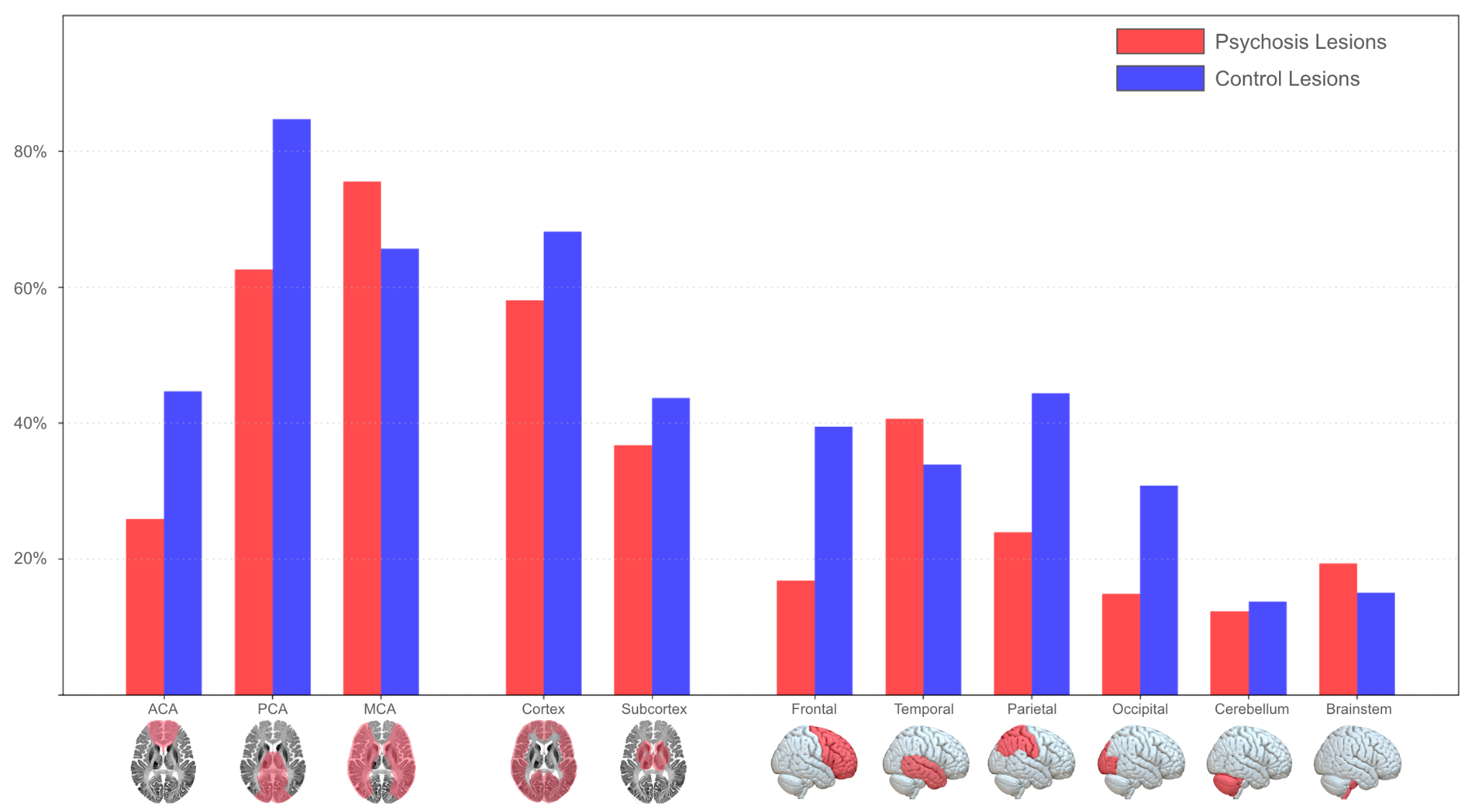

Distribution of lesions to cortex, subcortex, lobes, and vascular territories; illustrated as percentage of total lesions with voxels within each region. ACA=Anterior Cerebral Artery; PCA=Posterior Cerebral Artery; MCA=Middle Cerebral Artery.

**eFigure 4**

Voxel-wise two-sample t-test between lesions that cause psychosis and control lesions not associated with psychosis.

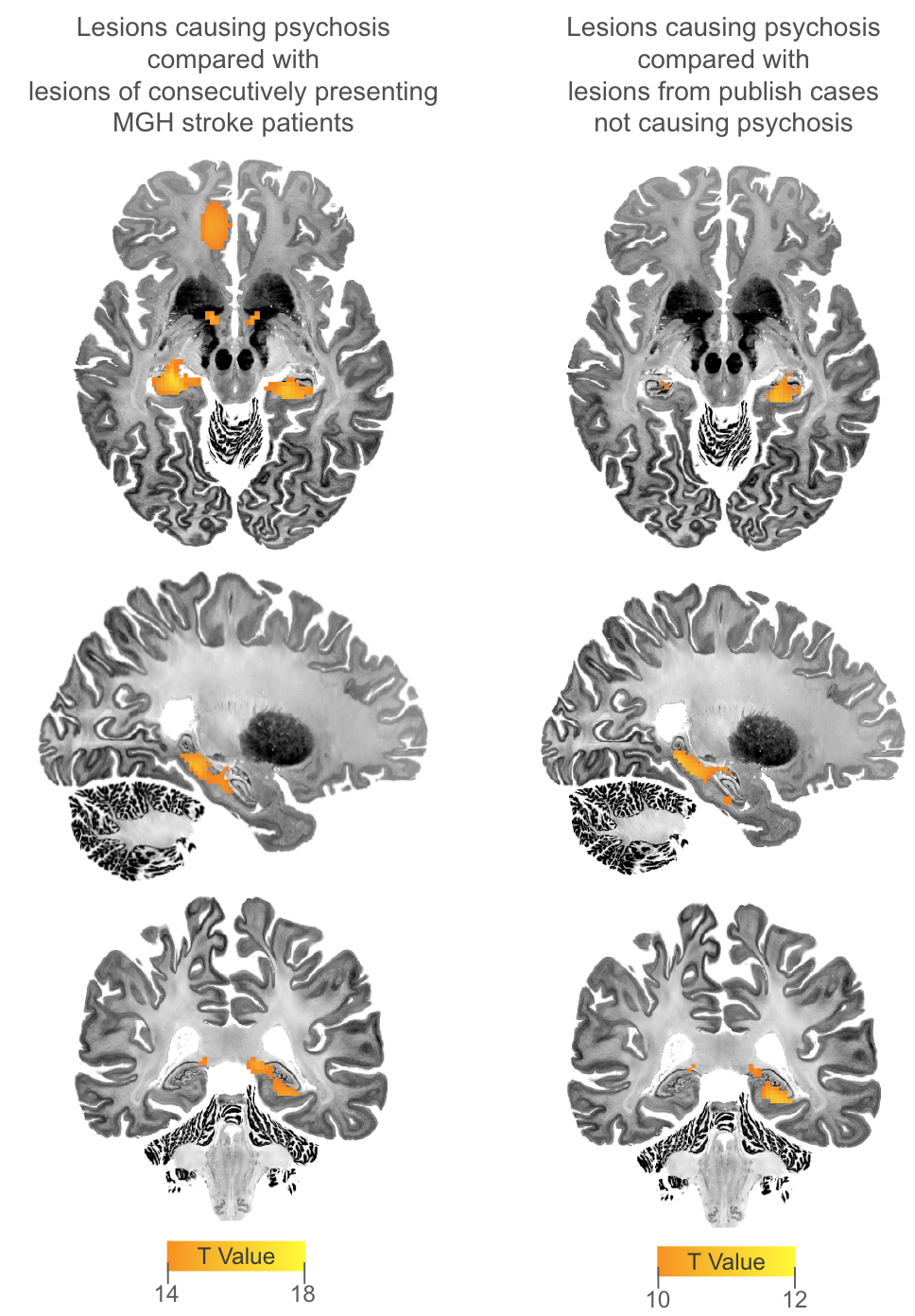

**eFigure 5**

Secondary Peak Regions of Psychosis Circuit

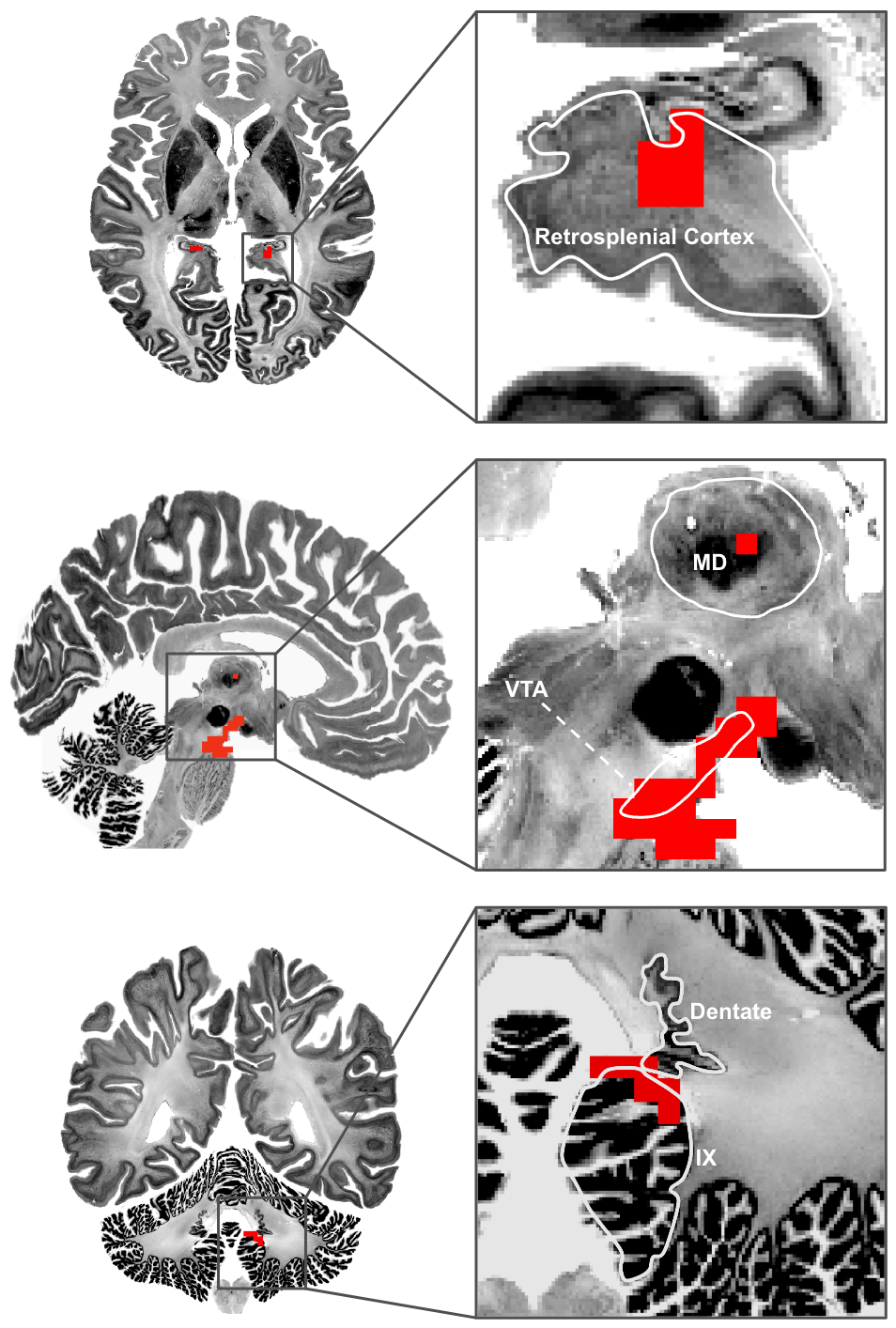

Convergence map depicting regions sensitive (overlap of functional connectivity of >75% of lesions) and specific (two-sample t-test (p_FWE_<1 x 10^-4^) comparing functional connectivity of lesions that cause psychosis and control lesions) to a circuit affected by lesions that cause psychosis. MD=Mediodorsal Nucleus of the Thalamus; VTA=Ventral Tegmental Area. Dentate=Dentate Nucleus of the cerebellum; IX=Lobule IX of the cerebellum

eFigure 6
Convergence of Sensitivity and Specificity Tests, Accounting for Age and Sex Covariates

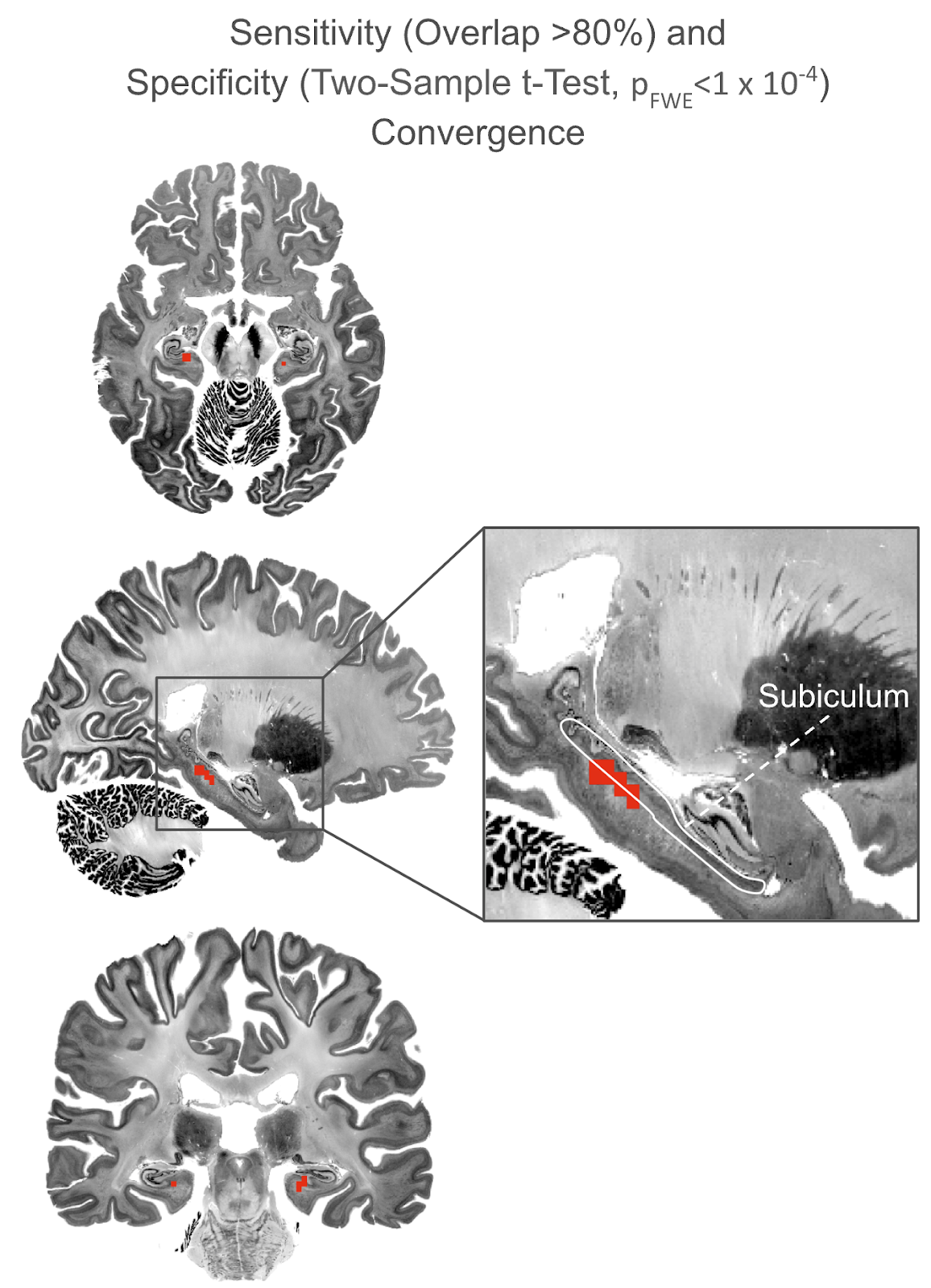

Sensitivity (Overlap>80%). Specificity (Voxel-wise Two-Sample t-Test, p_fwe_<1 x 10^-4^).

**eFigure 7**

Functional Connectivity of Lesions that Cause Psychosis is Consistent When Excluding Hippocampal Lesions

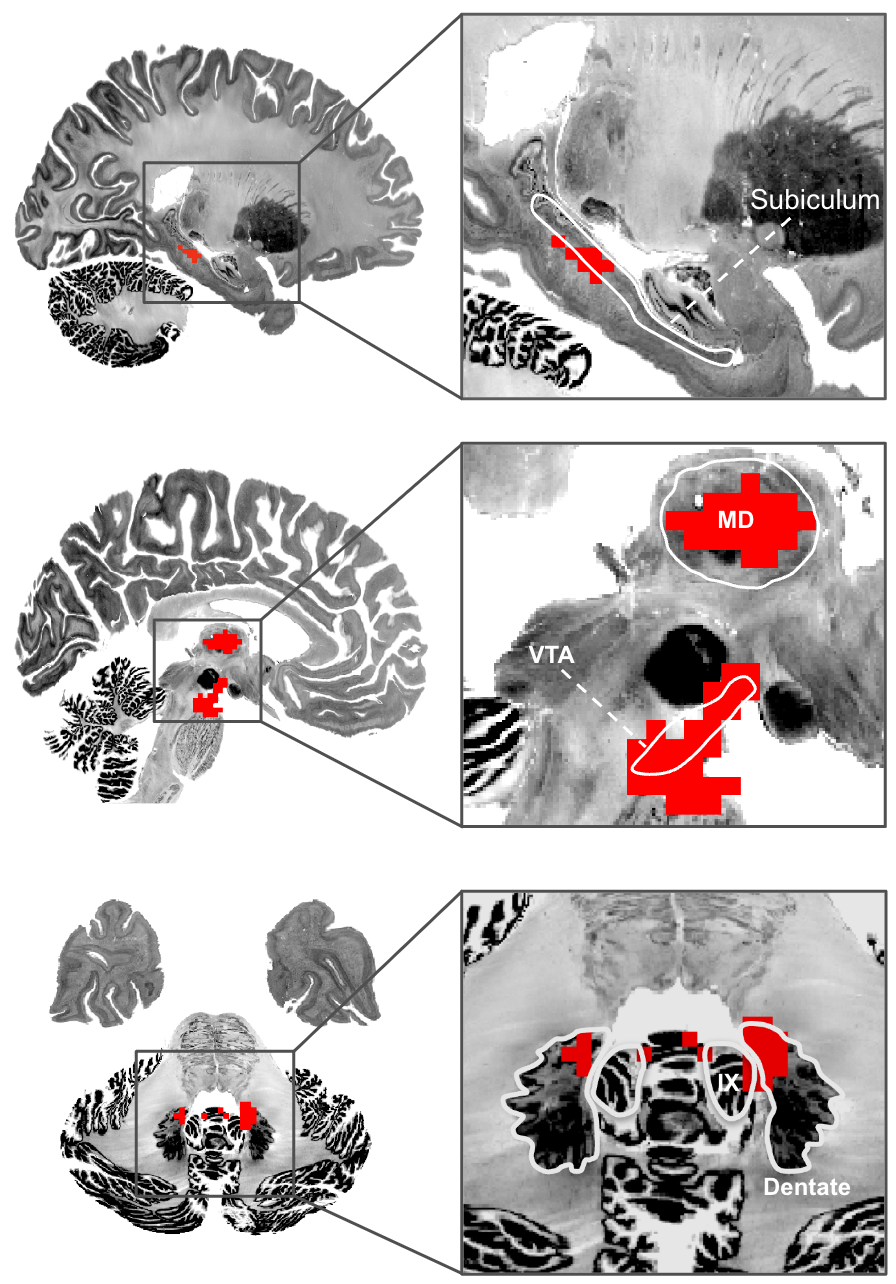

Convergence map depicting regions that are sensitive (overlap of functional connectivity of >75%) and specific (voxel-wise two-sample t-test against controls, p_FWE_<5 x 10^-5^) to a circuit affected by lesions that cause psychosis and do not directly intersect the hippocampus (n=98). Regions defined by the CoBrALab Merged Atlas^81^ and the Allen Brain Atlas^82^. MD=Mediodorsal Nucleus of the Thalamus; VTA=Ventral Tegmental Area. Dentate=Dentate Nucleus of the cerebellum; IX=Lobule IX of the cerebellum

**eFigure 8**

Leave-one-out analysis, leaving out the functional map of each lesion

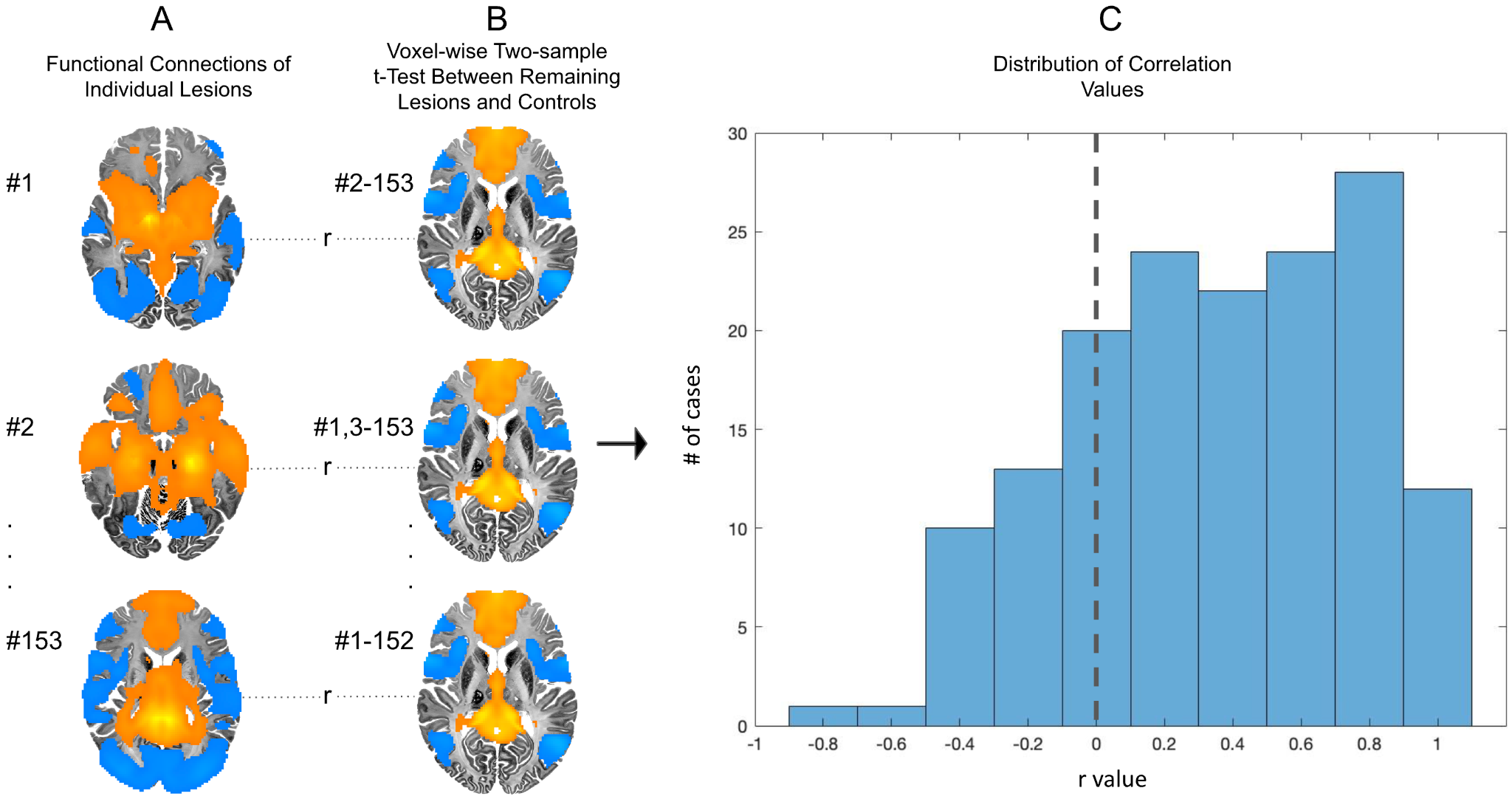

**a)**The functional connections of each lesion **b)** a voxel-wise two-sample t-test between the remaining 152 lesions and the control group **c)** a histogram displaying the distribution of spatial correlation values between the functional connections of each lesion and the specificity map generated from the functional connections of the remaining 152 lesions.

**eFigure 9**

Lesions that cause thought disorder map to different regions of a similar circuit.

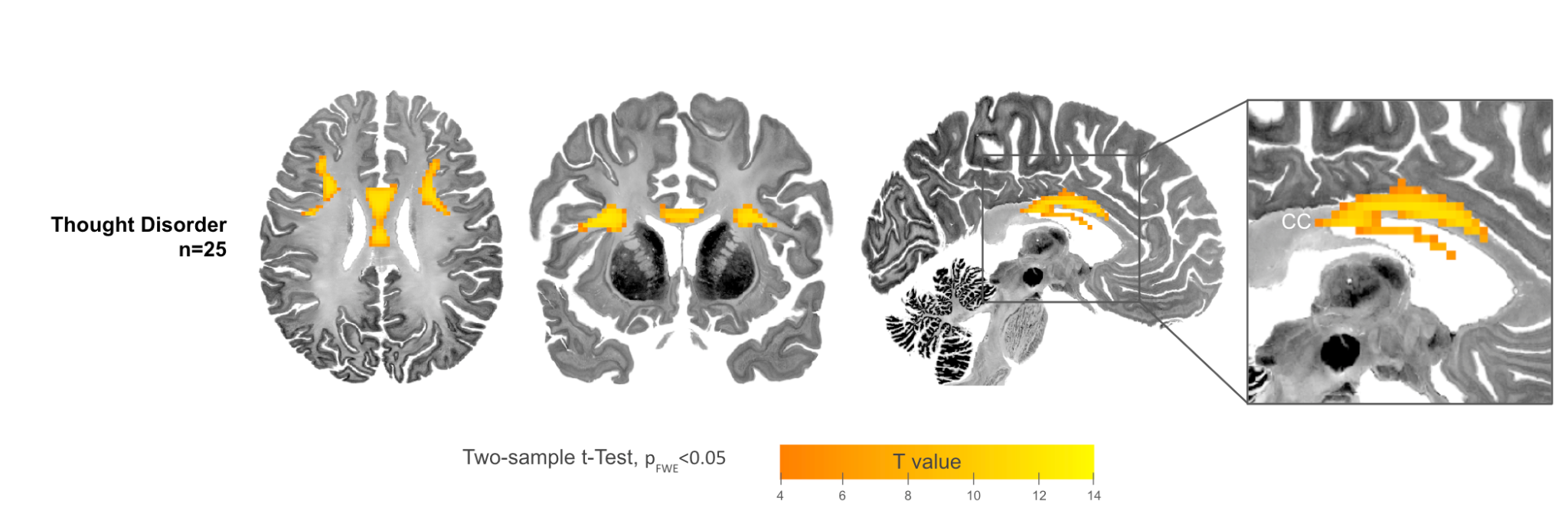

Voxel-wise two-sample t-tests between functional networks of lesions that cause thought disorder and lesions that cause other symptoms of psychosis (p_FWE_<0.05). CC= Corpus Callosum

**eFigure 10**

Lesions that cause isolated symptoms of psychosis map to distinct regions.

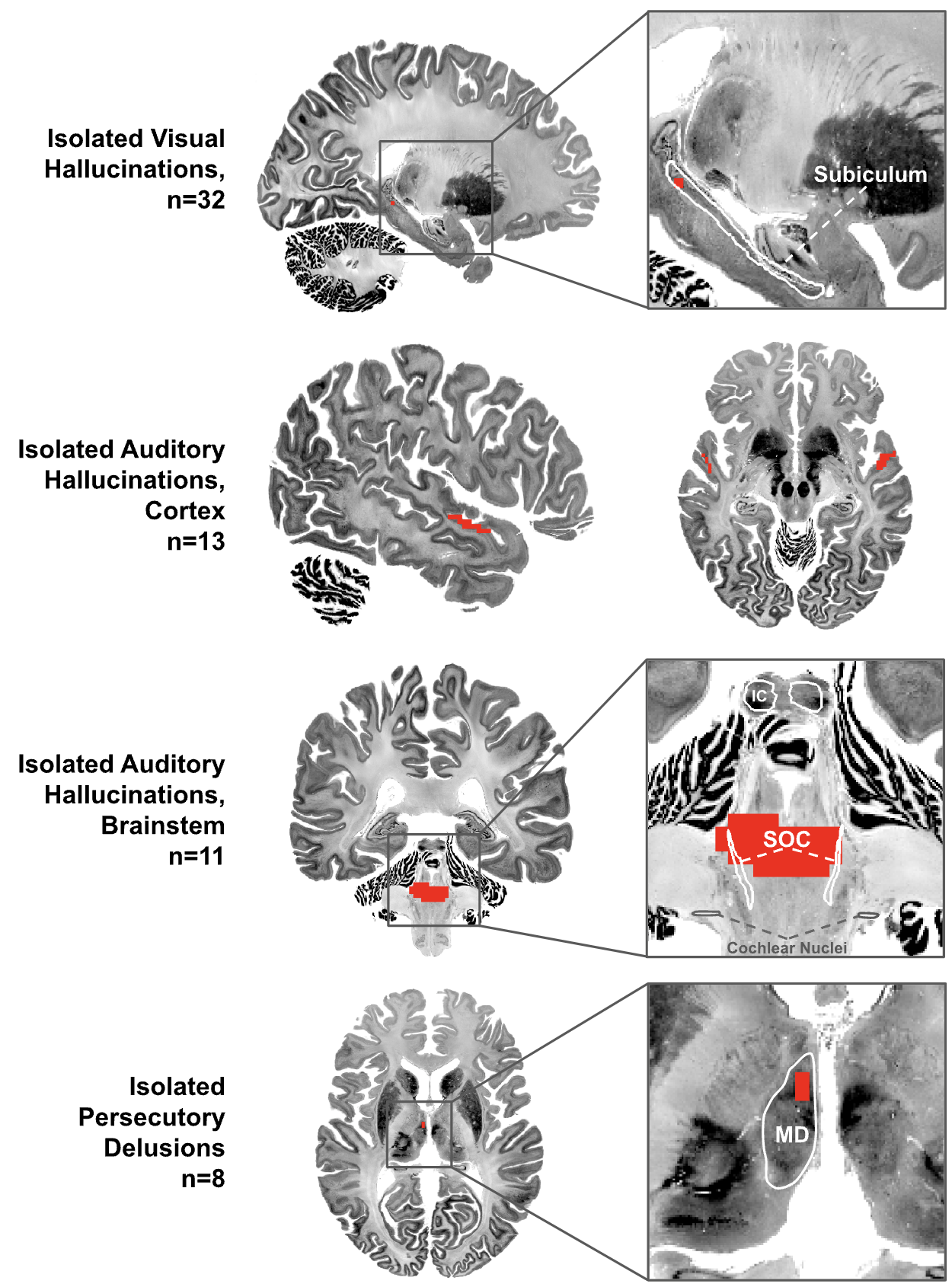

Conjunction of functional network overlap (>85%) and voxel-wise two-sample t-tests between each symptom group and controls (p_FWE_<0.05). Isolated visual hallucinations localize to the subiculum; isolated auditory hallucinations localize to the superior temporal gyrus or structures and lateral lemniscus white matter pathways of the subcortical auditory system^83^; isolated delusions localize to the right mediodorsal nucleus of the thalamus. IC=Inferior Colliculus; SOC=Superior Olivary Complex; MD=Mediodorsal Nucleus of the Thalamus

**eFigure 11**
Voxel-wise two-sample t-tests comparing lesions that cause psychosis with amnesia and lesions that cause only amnesia.

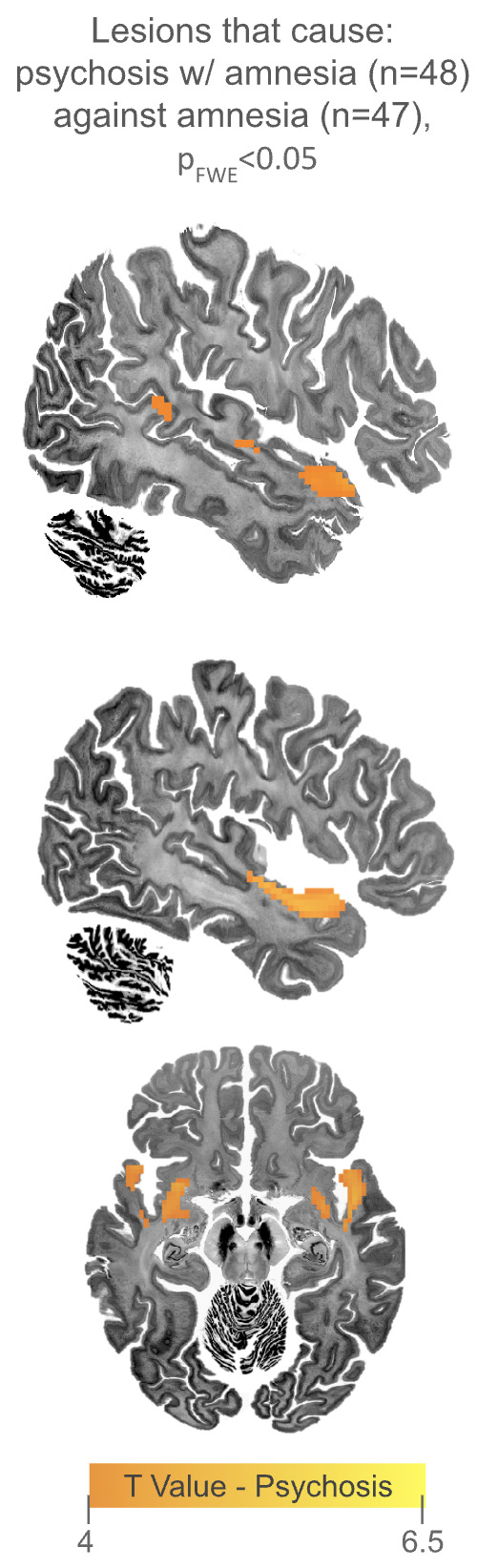

Voxel-wise two-sample t-test (p_FWE_<0.05) comparing connectivity maps of lesions that cause psychosis and amnesia and lesions that cause just amnesia. Lesions that cause psychosis and amnesia are more functionally connected to aspects of the superior temporal gyrus, the ventral claustrum, and part of the uncinate fasciculus, compared with lesions that cause just amnesia.
